# Multi-ancestry fine-mapping improves precision to identify causal genes in transcriptome-wide association studies

**DOI:** 10.1101/2022.02.10.479993

**Authors:** Zeyun Lu, Shyamalika Gopalan, Dong Yuan, David V. Conti, Bogdan Pasaniuc, Alexander Gusev, Nicholas Mancuso

**Author notes:** corresponding author: Nicholas Mancuso. contributed equally (Zeyun Lu, Shyamalika Gopalan). Authors Emails: Zeyun Lu Shyamalika Gopalan Dong Yuan David V. Conti Bogdan Pasaniuc Alexander Gusev.

## Abstract

Transcriptome-wide association studies (TWAS) are a powerful approach to identify genes whose expression associates with complex disease risk. However, non-causal genes can exhibit association signals due to confounding by linkage disequilibrium patterns (LD) and eQTL pleiotropy at genomic risk regions which necessitates fine-mapping of TWAS signals. Here, we present MA-FOCUS, a multi-ancestry framework for the improved identification of genes underlying traits of interest. We demonstrate that by leveraging differences in ancestry-specific patterns of LD and eQTL signals, MA-FOCUS consistently outperforms single-ancestry fine-mapping approaches with equivalent total sample size across multiple metrics. We perform 15 blood trait TWAS using genome-wide summary statistics (average N_EA_=511k, N_AA_=13k) and lymphoblastoid cell line eQTL data from cohorts of primarily European and African continental ancestries. We recapitulate evidence demonstrating shared genetic architectures for eQTL and blood traits between the two ancestry groups and observe that gene-level effects correlate 20% more strongly across ancestries compared with SNP-level effects. We perform fine-mapping using MA-FOCUS and find evidence that genes at TWAS risk regions are more likely to be shared across ancestries rather than ancestry-specific. Using multiple lines of evidence to validate our findings, we find gene sets produced by MA-FOCUS are more enriched in hematopoietic categories compared to alternative approaches (*P* = 1.73 × 10^−16^). Our work demonstrates that including, and appropriately accounting for, genetic diversity can drive deeper insights into the genetic architecture of complex traits.

## Introduction

Genome-wide association studies (GWAS) have identified genomic risk regions for numerous complex traits and diseases but leave unclear the underlying causal mechanisms responsible for risk. Multiple lines of evidence have suggested that genomic risk is imparted through perturbed regulation of nearby target genes, which predicts that the steady-state abundance of expression levels at target genes is associated with disease risk^1-6^. Transcriptome-wide association studies (TWAS)^1,2^, which explicitly test this hypothesis, have been successful in identifying novel genomic risk regions and specific genes that influence complex diseases^7-9^. Much of TWAS’ recent success is due to the use of genetically predicted, rather than directly assayed, gene expression, which enables its application to existing large-scale GWAS, thus greatly increasing statistical power. Recently, we and others have demonstrated that TWAS also suffer from confounding due to eQTL pleiotropy and LD, which can induce correlation in test statistics between causal and non-causal genes in an analogous manner to causal and tagging variants in GWAS^10-16^.

Despite these recent breakthroughs, our understanding of the genetic architecture of complex traits has been limited by a lack of diversity in human genetics studies: individuals with primarily European genetic ancestry comprise 79% of all GWAS participants, despite representing only 16% of the global population^17^. Although risk loci frequently replicate across ancestries^18-22^, the linkage disequilibrium (LD) patterns, minor allele frequencies (MAF), and the number of causal variants and their effect sizes can vary across genetic ancestries^21^. This heterogeneity in genetic architecture hinders clinical applications of GWAS such as polygenic risk scores (PRS), an issue that has been highlighted by the poor portability of PRS models across ancestries^23,24^. On the other hand, recent trans-ancestry design of GWAS have highlighted the benefits of taking an integrative, multi-ancestry approach to studying complex disease biology, both by leveraging genetic heterogeneity across human groups to aid in fine-mapping, and by enabling the discovery of ancestry-specific disease etiologies^20,21,25-27^. As with GWAS, we expect the integration of genetically diverse datasets into TWAS methodologies will improve our understanding of trait architectures that are both shared and unique to particular genetic ancestries^28,30,32^.

In this work, we present MA-FOCUS (Multi-Ancestry Fine-mapping Of CaUsal gene Sets), an approach that integrates GWAS, expression quantitative trait loci (eQTL), and LD data from multiple ancestries to assign a posterior inclusion probability (PIP) that a given gene explains the TWAS signals at a risk region^29,31,33^. It uses inferred PIPs to compute credible sets of causal genes at a predefined confidence level *ρ* (**Figure 1**). A key feature of MA-FOCUS is that it does not assume that the eQTL architecture underlying gene expression is shared across ancestries^34,35^. Instead, MA-FOCUS assumes only that the causal genes for a focal trait or disease are shared across ancestries. It is expected that gene-level effects are likely more transferable across ancestry groups than SNP-level effects as genes are inherently a more meaningful biological unit^36^. As a result, MA-FOCUS leverages cross-ancestry heterogeneity in LD patterns and eQTL associations to identify causal genes with improved precision and accuracy when compared with alternative approaches.

**Figure 1.**
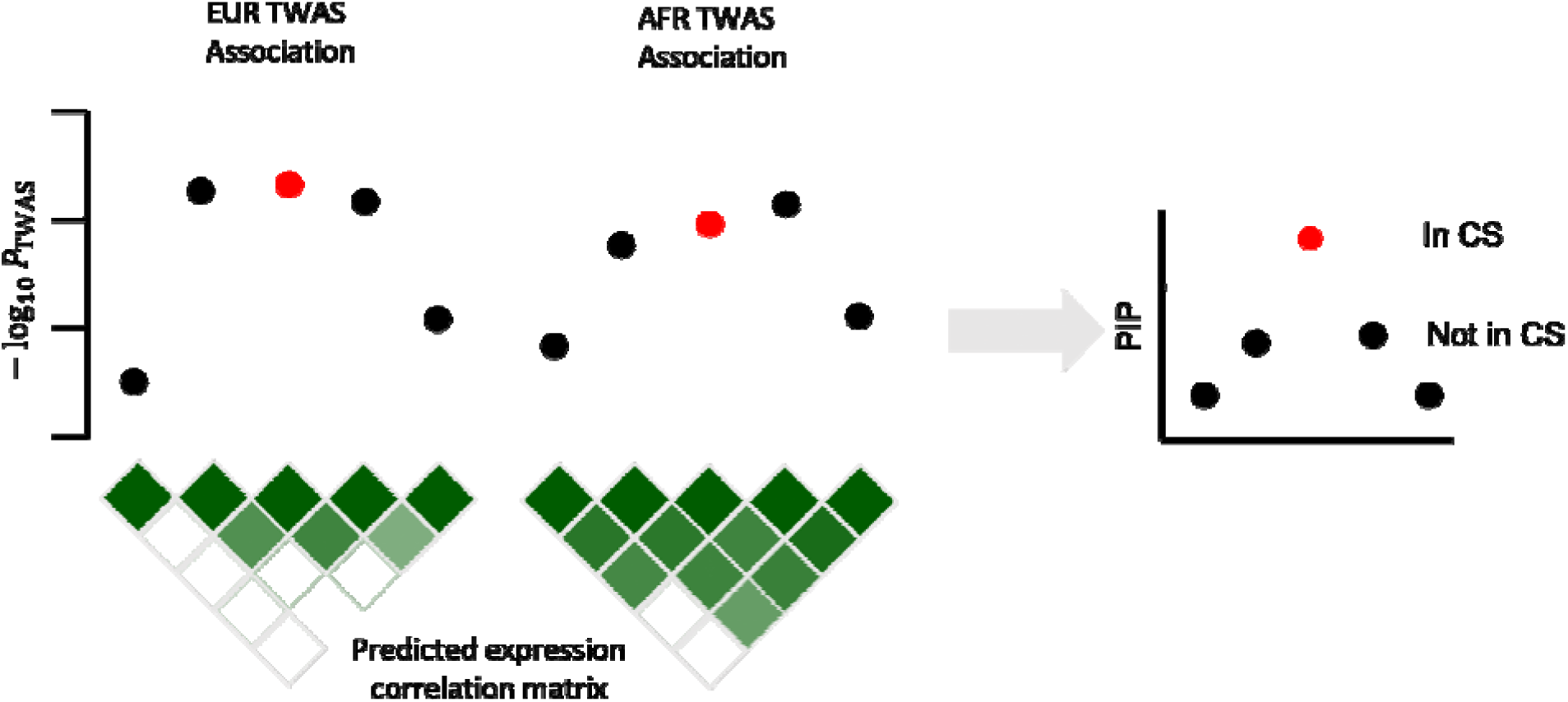
Example of correlated TWAS associations at a shared risk region. Toy example of TWAS Manhattan plots for EUR and AFR ancestries illustrating association signals at a locus for the causal gene (in red) and tagging genes (in black). The correlation among association signals is a combined result of eQTL signals and linkage disequilibrium (LD; see Methods). By accounting for heterogeneity in eQTL effect sizes and LD across different ancestries, MA-FOCUS produces smaller gene credible set with more posterior probability assigned to the causal gene.

By performing extensive simulations, we demonstrate that MA-FOCUS consistently outperforms the analogous single-ancestry method with equivalent total sample size, as well as a ‘baseline’ approach based on meta-analyzed GWAS statistics from different ancestries^37,38^. In addition, we show that MA-FOCUS is robust in simulations where the trait-relevant tissue is missing, and a proxy tissue is used instead. To illustrate its applicability on real multi-ancestry data, we conduct multiple TWAS and fine-mapping analyses with MA-FOCUS for 15 blood traits in European and African ancestry cohorts using large-scale GWAS summary statistics^18^ (average N_EA_=511k, N_AA_=13k) and eQTL weights calculated from the Genetic Epidemiology Network of Arteriopathy (GENOA) study^39^ (N_EA_=373, N_AA_=441). We recapitulate results demonstrating the shared genetic architecture for gene expression and blood traits between the two ancestries. We also find evidence that gene-level effects inferred from TWAS correlate 20% more strongly across ancestries when compared with SNP-level effects. Fine-mapping the 22 genomic regions that contain TWAS signals for both ancestry groups, we find MA-FOCUS identifies genes relevant to hematopoietic and cardiovascular disease etiology that are missed by the baseline approach. Using multiple validation strategies^40^, we show genes in MA-FOCUS credible sets are more strongly enriched for hematological measurement categories (meta-analysis P-value of 1.73 × 10^−16^ compared to 2.91 × 10^−11^) compared to the baseline approach. Overall, our analyses using MA-FOCUS emphasize the importance of incorporating genetic information from diverse genetic ancestries to drive new insights into the genetic architecture of complex traits.

## Materials and methods

### Multi-ancestry FOCUS Model

For the *i*^*th*^ of *k* total ancestries, we model a centered and standardized complex trait 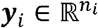 from *n*_*i*_ individuals as a linear combination of gene expression levels 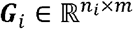 at *m* genes as

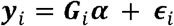

where ***α*** ∈ ℝ^*m*^ are the causal effects of gene expression on the complex trait, which are shared across all ancestry groups, and 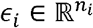 is random environmental noise with 𝔼(***ϵ***_*i*_) = 0 and 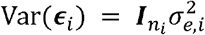. Additionally, we model ancestry-specific gene expression as a linear combination of genotypes ***X***_*i*_ as

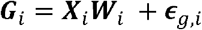

where 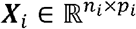 is the centered and standardized genotype matrix at ***p***_*i*_ single-nucleotide polymorphisms (SNPs), 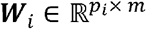 is the ancestry-specific eQTL effect-size matrix, and 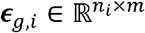 is random environmental noise.

Performing a TWAS using predicted gene expression requires the latent ancestry-matched eQTL weights ***W***_*i*_ which are unknown. In practice, we use expression weights ***Ω***_*i*_ estimated from an independent, ancestry-matched eQTL reference panel using penalized linear models (or Bayesian counterparts)^1,2^. We model the *i*^*th*^ ancestry’s marginal TWAS summary statistics for the gene *j* with complex trait ***y***_*i*_ as 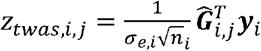 where 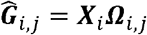 is the predicted expression imputed by the eQTL panel. By algebraic expansion for *m* genes, we have

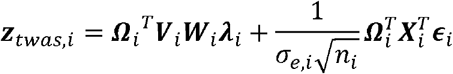

where we re-parameterize the causal effects of gene expression as 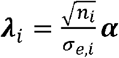 and the ancestry-matched LD at *p*_*i*_ SNPs as 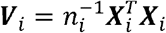. Assuming expression weights ***Ω***_*i*_, and causal effects ***α*** are fixed, we can compute the sampling distribution of ***z***_*twas,i*_ as

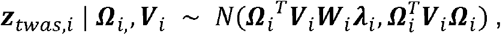

and as sample size increases, 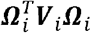 asymptotically approaches *Ω*_*i*_^*T*^*V*_*i*_*W*_*i*_.

Next, we model a prior for the causal effects as 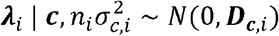 where 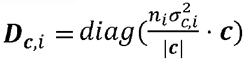, *c* is an *m* × 1 causal configuration binary vector where *c*_*j*_ = 1 if the *j*^*th*^ gene at KI the region is causal (0 otherwise), |***c***| denotes the length of ***c***, and 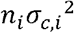 denotes the samplesize scaled causal effect prior variance^10^. We marginalize ***λ***_*i*_ out to obtain the TWAS sampling distribution conditioned on a causal gene set as

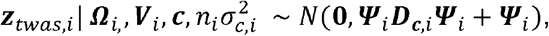

where 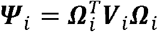 is the estimated expression correlation matrix. We assume that the causal genes underlying a complex trait are shared across ancestries, which we model by sharing the *c* vector across ancestries. Since we do not know the causal genes indicated by ***c*** beforehand, we adopt a Bayesian approach and compute the posterior for a given causal configuration *c* as

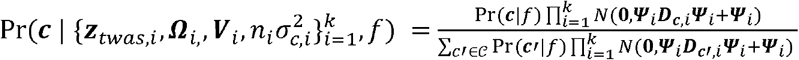

where Pr(***c***|*f*) = *f* ^|*c*|^(1−*f*)^(*m*−|*c*|)^ for some prior causal probability *f* and *𝒞* is the space of causal gene configurations. In practice we set *f* to be 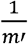 where *m*’ ≥ *m* denotes the number of known, and not necessarily tested, genes at the region. Intuitively, this reflects the naive expectation that a given risk locus contains a single causal gene. For computational tractability, we constrain the space defined by *𝒞* to exclude complex configurations with |***c***| > 3. In addition, our likelihood, and thus posterior, depends on 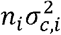 which governs the variance of scaled causal gene effects ***λ***_*i*_. Previously, we recommended using a genome-wide mean 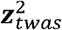 as a heuristic, which works well under polygenic architectures^10^, but may perform poorly in sparser situations. Motivated by ref^41^, here we describe a local heuristic that estimates 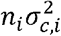 as

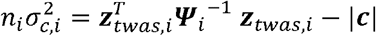

which is an unbiased estimator of causal effect variance ***α***^*T*^***Ψ***_*i*_***α*** (see **Appendix**). In the case of negative estimates, we instead use 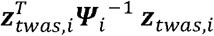.

### Computing Posterior Inclusion Probabilities and *ρ*-Credible Sets

Our model describes the posterior probability for a given causal configuration ***c*** across ancestries; however, we are more interested in the probability that a specific gene is causal across ancestries. We define the posterior inclusion probability (PIP) for the *j*^*th*^ gene by marginalizing over all causal configurations *c* where *c*_*j*_ = 1 as:

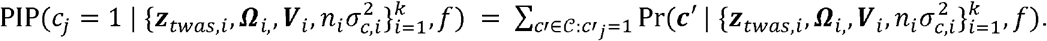

To capture the probability that none of the genes included in our analysis explain the observed TWAS Z-scores at a risk region, we include the null model as a possible outcome in the credible set, 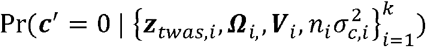. To compute a *ρ*-credible set^29,31,33^, where *ρ* reflects the desired confidence that a gene set contains a causal gene, we take a greedy approach that traverses genes ordered decreasingly by their locus-normalized PIPs until the cumulative sum reaches at least *ρ*.

### Overview of the simulation pipeline

Here we provide a high-level summary of our multi-ancestry TWAS simulation pipeline, which is described by five main steps (**Figure S1**), with details for each step described in the following sections. First, we computed approximately independent LD blocks and sampled genotypes for GWAS and eQTL reference panels in two ancestry groups. Second, we simulated ancestry-matched eQTL data using simulated eQTL reference genotypes from the first step, sampled eQTL effects under a sparse architecture, and simulated gene expression at causal and non-causal genes. Third, we simulated a complex trait in the ancestry-matched GWAS data as a linear function of eQTL effects of the causal gene from the second step and simulated GWAS genotypes from the first step. Fourth, we performed ancestry-matched TWAS using penalized models fitted in the respective eQTL reference panels. Fifth, we performed fine-mapping using single-ancestry FOCUS and MA-FOCUS. We provide details for each step below.

### Computing independent LD blocks and simulating reference eQTL panels

We performed simulations using genotype data from phase 3 of the 1000 Genomes Project for individuals of European (EUR; N=490) and African (AFR; N=639) ancestries (see **Table S1**)^42^. We restricted genotypes to high-quality HapMap SNPs and filtered for missingness (> 1%), minor allele frequency (MAF < 1%), and violations of Hardy-Weinberg equilibrium (HWE mid-adjusted P-value < 1e-5). To identify approximately independent regions that are consistent with both EUR and AFR ancestries, we used a recently described extension of LDetect that considers LD information from multiple ancestries^21,43^. Briefly, we constructed chromosome-wide ancestry-matched LD matrices ***V***_*i*_ and computed a chromosome-wide trans-ancestry LD matrix ***V***_*trans*_ such that it incorporates shared recombination loci across ancestries (see ref^21^). Applying LDetect^21,43^ to ***V***_*trans*_ resulted in 1278 approximately independent LD blocks. We sampled 100 blocks that carried between 5 and 8 annotated genes (based on hg19 RefSeq release 63) as risk regions. Additionally, we extended each LD block 500kb upstream of the first gene’s transcription start site (TSS) and 500kb downstream of the last gene’s transcription end site (TES) and updated ***V***_*i*_ accordingly.

At each risk region, we simulated 10 genes whose expression is under partial genetic control by first sampling the number of eQTLs for the *j*^*th*^ gene, *k*_*j*_ = max(1, Poisson(2)). Next, we assigned *k*_*j*_ SNPs uniformly at random to be eQTLs (out of *p* total for a given locus) and simulated *p* × 1 effect-sizes vector 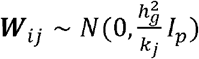 at the *k*_*j*_ causal eQTLs and 0 at the *p* − *k*_*j*_ non-causal SNPs where 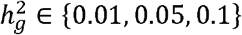 is the proportion of variance in gene expression attributable to *cis*-eQTLs (i.e. SNP heritability of gene expression)^5,44^. In addition, we simulated eQTLs as either independent or shared across ancestries; in the former case, SNPs and their effect sizes were chosen for each ancestry individually (under shared 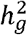 and *k* parameters) as described above, while in the latter, these were chosen once and then fixed for all ancestries^34,35^. Then, we simulated a *n*_*i,eQTL*_ × *p* centered and standardized continuous genotype matrix ***X***_*i,eQTL*_ using a multivariate normal distribution *N*(0, ***V***_*i*_) where *n*_*i,eQTL*_ is the ancestry-matched eQTL panel sample size. For gene *j*, we calculated expression ***G***_*ij*_ according to ***G***_*ij*_ = ***X***_*i,eQTL*_***W***_*ij*_ + ***ϵ***_*g,ij*_, where 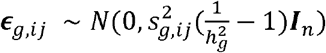 is random environmental noise for expression ***G***_*ij*_, and 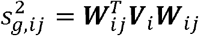. To estimate ancestry-matched expression weights **Ω**_*ij*_, we regressed ***G***_*ij*_ on ***X***_*i,eQTL*_ using least absolute shrinkage and selection operator (LASSO) regularization. To simulate eQTL effects when only a genetically correlated proxy tissue is available, we sampled proxy eQTL effects 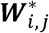 under a bivariate normal distribution as

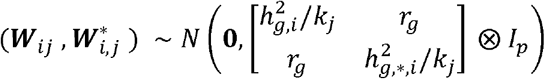

where ***W***_*ij*_ are the causal tissue eQTLs and *r*_*g*_ ∈ {0,0.3,0.6,0.9,1} is the genetic covariance between two tissues.

### Simulating complex traits and statistical fine-mapping of TWAS

To reflect the practical reality that participants in GWAS and eQTL panels are usually different, we re-simulated genotypes ***X***_*gwas,i*_ ∼ *N*(0,***V***_*i*_) at the risk region to compute GWAS summary statistics while keeping eQTLs ***W***_*ij*_ of the 10 simulated genes from the previous step. Then, we randomly sampled one gene as causal and used its eQTLs to simulate complex trait ***y***_*i*_ as

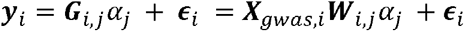

where *α*_*j*_ ∼ *N*(0,1) is the causal gene expression effect, 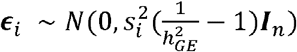 is random environmental noise for complex trait ***y***_*i*_ where 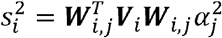, and 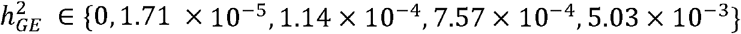 is the proportion of complex trait variation explained by the genetic component of gene expression. Next, to compute ancestry-matched GWAS summary statistics, we performed linear regression on the complex trait ***y***_*i*_ marginally for each SNP in ***X***_*gwas,i*_ and calculated GWAS Z-scores ***z***_*gwas,i*_ using the resulting Wald test statistic. We then performed an ancestry-matched summary-based TWAS using predicted expression ***z***_*twas,i*_ for each gene with 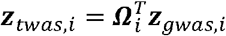.

Lastly, we performed TWAS fine-mapping using single-ancestry FOCUS^10^ and multi-ancestry MA-FOCUS on ***z***_*twas,i*_ to generate 90% credible sets for each ancestry and under the joint model, respectively. To determine whether the improvement of MA-FOCUS is solely due to increased sample sizes, we also evaluated a ‘baseline’ approach^37^. Specifically, the baseline approach consists of computing meta-analyzed GWAS statistics as 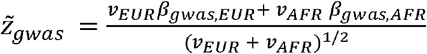, where 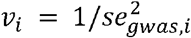 is the inverse variance weight. Rather than constructing meta-analysis expression weights, 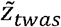 is then computed by using the EUR expression weights ***Ω***_*EUR*_. Finally, fine-mapping is conducted on 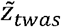 using single-ancestry FOCUS and 90% credible sets are computed. In all, we ran four methods (EUR FOCUS, AFR FOCUS, baseline, and MA-FOCUS) on 100 LD blocks to output one credible set per LD block per method. To test whether including information from additional ancestries of diverse genetic ancestries increases the performance of MA-FOCUS, we evaluated scenarios that also include individuals simulated using 1000G East Asian (EAS; N=481) ancestry data^42^ (**Table S1**), and performed MA-FOCUS on three ancestries by fixing per-ancestry eQTL sample size, 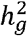, and 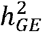 and allowing the total GWAS sample size to vary.

### Simulating ancestry-specific genetic architectures and data-missing cases

A central assumption of MA-FOCUS is that different ancestries share the same causal genes and their effect sizes. To characterize the performance of MA-FOCUS in cases where this assumption is partially violated, we simulated cases where the mediating gene-trait effects 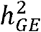 are ancestry-specific as a heuristic to represent heterogeneity in genetic architectures. Additionally, in practice, eQTL panels for a particular tissue of interest may be either unavailable or underpowered due to small sample size. To evaluate the performance of MA-FOCUS in cases where relevant eQTL data are unavailable^45^, we tested two scenarios that used different types of “proxy” data^1,8,10,34,35,46^. First, we simulated cases where the trait-relevant tissue was unavailable in AFR, and a proxy tissue from the same ancestry with correlated gene expression was substituted. Second, we simulated cases where eQTL weights for AFR were entirely unavailable, and weights from EUR were used instead. The latter differs from the baseline approach in that the TWAS and FOCUS were conducted with ancestry-matched, not meta-analyzed, GWAS results.

### Description of simulation parameters and fine-mapping performance metrics

We compared MA-FOCUS results to single-ancestry FOCUS results for EUR and AFR, and the baseline approach across multiple simulations, which varied according to whether eQTLs were shared or not. We also varied four additional parameters: GWAS sample sizes, eQTL panel sample sizes, *cis*-SNP heritability of gene expression 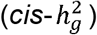, and the proportion of trait variance explained by genetically-regulated gene expression 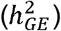. Unless stated otherwise, the simulation parameters were set to defaults of 100,000 for the per-ancestry GWAS sample size, 200 for the per-ancestry eQTL panel size, expression 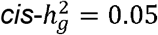 and trait 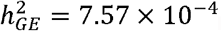. We evaluated fine-mapping performance based on three metrics: mean PIP of the causal genes, mean 90% credible set size, and frequency in which the causal genes are included in 100 90% credible sets per simulation (sensitivity). We fit linear regression adjusted for corresponding parameters to report one-sided Wald test P-value.

### Fitting SNP-based prediction models of LCL expression in the GENOA study

To calculate ancestry-specific gene expression weights in real data, we used genotype and lymphoblastoid cell line (LCL) derived gene expression data from European ancestry (EA) and admixed African American (AA) individuals from the GENOA study^39^. Genotype data were generated using Affymetrix and Illumina genotyping arrays; in total, 1,384 EA and 1,263 AA individuals were assayed on the Affymetrix 6.0 array, 20 EA and 269 AA on the Illumina 1M array, and 103 EA on the Illumina 660k array. All genotype data analyses were conducted using PLINK 1.9, vcftools, and bcftools^47-50^. We imputed genotype data using the TOPMed server, implementing minimac4 v1.0.2 and eagle v2.4 phasing^51^. Each ancestry dataset was imputed separately, except for EA individuals assayed on Illumina arrays, which we merged prior to imputation. We retained bi-allelic SNPs with good imputation quality (*r*^2^ > 0.6) for both EA and AA cohorts, filtered for MAF < 1% and for HWE P-value < 1 × 10^−6^ resulting in 1,160,917 and 1,330,340 quality-controlled (QC) SNPs for EA and AA, respectively. We used GCTA^52^ to compute genotype principal components (PCs) and genetic relatedness matrices within the EA and AA cohorts after further filtering for SNPs with imputation *r*^2^ > 0.9 and low pairwise LD (using --indep-pairwise 200 1 0.3 in PLINK^48^). For computing genotype PCs, we filtered out individuals such that no pair exhibited a relatedness coefficient greater than 0.05, resulting in 373 EA and 441 AA individuals. For downstream eQTL model fitting, we used only HapMap SNPs^52^.

Expression data for the EA and AA cohorts were assayed at 16,944 and 32,881 genes (overlap of 14,797) on the Affymetrix Human Exon 1.0 and Affymetrix Human Transcriptome 2.0 arrays, respectively, and processed by Shang et al^39^. In this context, genes refer to any regions that express RNA, and not necessarily the ones that have protein-coding function. After lifting over the expression data to GRCh38, for each gene in its respective ancestry, we ran FUSION^1,7,9^ to estimate genetic variance 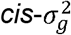 and 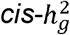, and to calculate ancestry-specific eQTL weights, limiting the analysis to SNPs falling within a window including 500kb up and downstream of each gene’s transcription start-site and transcription end-site, as defined in hg19 RefSeq release 63. We included 30 gene expression PCs, 5 genotype PCs, age, sex, and genotyping platform as covariates in building these SNP models^1,7,9^. We identified 3,680 and 4,291 genes in EA and AA, respectively, with an estimated 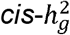 of at least 0.01 (nominal p-value < 0.01) of which 2,496 genes overlapped both ancestries. We limited our downstream analyses to 4,646 unique genes that had evidence for significant 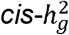, as defined above, in either ancestry and non-zero weights in both ancestries^1,7,9^.

### Validation of LCL prediction models in GEUVADIS

To validate our estimated ancestry-specific gene expression weights derived from the GENOA study^39^, we obtained paired genotypes and LCL-derived mRNA expression data at 22,721 genes for 373 EUR participants and 89 Yoruba in Ibadan (YRI) participants from the GEUVADIS study^53^. We performed the same relatedness- and variant-based filtering as described above, which resulted in 358 EUR and 89 YRI participants and 8,403,216 and 14,855,241 SNPs, respectively. Restricting to the 4,581 genes that overlapped with GENOA, we estimated 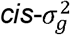 and 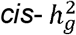 of LCL gene expression in GEUVADIS using the Genome-based restricted maximum likelihood (GREML) approach, with genotypes limited to 500kb up and downstream window around each gene as described above, and adjusted for participants’ sex and 3 genotype PCs^52^. Next, we predicted LCL expression for GEUVADIS participants using GENOA-based expression weights and calculated the coefficients of determination (*r*^2^) between predicted and measured expression levels.

### TWAS and fine-mapping of 15 blood traits from GWAS summary data

We obtained published GWAS summary statistics for 15 blood traits (**Table S2**) from Chen et al^18^. After lifting SNPs over to GRCh38 and updating their identifiers to dbSNP v153, we used LDSC munge^54^ to perform quality control filtering of summary statistics, based on imputation INFO scores > 0.9, MAF > 0.01, and chi-squared statistics < 80 to limit the influence of outlier SNPs. We flipped alleles as necessary for consistent orientation across European ancestry and African ancestry GWAS statistics. The average GWAS sample size was 511,471 for European and 13,298 for African ancestries across all SNPs and all 15 blood traits, reflecting ∼40-fold difference in sample sizes.

As in our simulations, we calculated TWAS *z* scores^1^ of EA, AA, and baseline approach for each trait by leveraging corresponding GWAS summary statistics^18^, FUSION-fitted LCL eQTL reference weights^1,39^, and estimated LD from 1000 Genomes^42^. To shed light on ancestry similarity in genetic architecture, we computed cross-ancestry correlation of GWAS and TWAS effect-size estimates for genome-wide SNPs and genes using a blocked jack-knife approach; to adjust for sample size differences, we normalized GWAS and TWAS effect sizes by dividing by square root of GWAS sample sizes. To compute an average across all 15 blood traits, we meta-analyzed individual correlations across 15 blood traits and tested the difference with pooled standard error. Next, we fine-mapped the original resulting TWAS Z-scores using MA-FOCUS, single-ancestry FOCUS, and the baseline approach, focusing on independent genomic regions which exhibited transcriptome-wide significant signals (*P* < 0.05/4579, the number of genes with TWAS statistics) in both EA-specific and AA-specific TWAS, and annotated them based on their inclusion in the 90% credible set, as described above. To provide evidence of the causal genes being shared, rather than ancestry-specific, we performed Bayesian model comparison and calculated log-Bayes factors for each gene in a MA-FOCUS credible set as

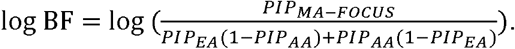

### Validation of blood trait fine-mapping results

To determine if the genes prioritized by MA-FOCUS are more biologically meaningful than those prioritized by other methods, we validated credible sets using four different approaches. First, we performed a gene set enrichment analysis for genes identified in credible sets (i.e. aggregating genes identified across all loci) for a given fine-mapping method and blood trait using the R package enrichR^55,56^. We manually selected 20 trait categories related to hematological measurement in DisGeNET, a database of curated gene-trait associations^40^, based on the most relevant body system using MeSH (see **Web resources**) and EFO^57^ ontology hierarchies (**Table S3**). We counted the number of significantly enriched categories with Bonferroni correction (*P* < 0.05/*n* where *n* is the number of enrichment testing) for each method and performed meta-analysis on these categories using Fisher’s Method. Second, we performed enrichment analyses comparing each blood-trait and fine-mapping specific gene set with the DisGeNET-curated gene set for the equivalent blood trait. Third, we evaluated gene sets using a previously published “silver standard” (see **Web resources**), to determine whether they better predict causal genes of 159 blood-related mendelian and rare diseases (**Table S4**). Since these diseases are monogenic or oligogenic, their causal genes are affirmative in high confidence and are likely to have moderate effects on blood-related complex traits. Leveraging database from Online Mendelian Inheritance in Man (OMIM) and Orphanet, we performed logistic regression to calculate areas under the ROC curve within each method, and each blood-related trait in Chen et al.

## Results

### Multi-Ancestry FOCUS improves power to identify causal genes in simulations

We first evaluated the performance of MA-FOCUS in simulations and compared it with the baseline approach, which consists of GWAS meta-analysis across ancestries followed by TWAS and fine-mapping with a single ancestry’s weights (see **Methods**). Briefly, we simulated a complex trait as a function of genetically-regulated gene expression for both ancestries when the causal tissue was known (see **Methods**) while varying GWAS and eQTL sample sizes and features of the underlying genetic architecture. Across all simulation scenarios where causal eQTLs were independent across ancestries, we found MA-FOCUS reported higher PIPs for causal genes than the baseline approach (0.62 compared with 0.45; *P* = 9.05 × 10^−40^), smaller credible sets (4.89 compared to 6.62; *P* = 2.13 × 10^−131^), and higher sensitivity (88.30% compared to 81.30%; *P* = 9.35 × 10^−9^). Specifically, consistent with previous TWAS and TWAS fine-mapping simulation studies^1,10^, we observed performance improved as GWAS and eQTL sample sizes increased, likely reflecting increased statistical power (**Figures 2, S2**). We found that increasing eQTL panel size affected MA-FOCUS sensitivity more dramatically than increasing GWAS sample size. For instance, increasing the eQTL panel size two-fold, from 200 to 400, improved sensitivity by 6% from 91% to 97% while a same proportionate increase in the GWAS sample size, from 100,000 to 200,000, only increased sensitivity by 2% to 93% (**Figure 2, S2**). We re-performed these simulations assuming that the causal eQTLs are shared across ancestries and observed that MA-FOCUS consistently outperformed the baseline (**Figure S3**). However, this performance advantage was slightly attenuated compared to the independent eQTL setting, highlighting the ability of MA-FOCUS to improve performance while being agnostic to eQTL architecture. Hereafter, we focus on presenting results where eQTLs were simulated independently in each ancestry to highlight MA-FOCUS’ potential advantage in real-world applications where eQTLs exhibit heterogeneity across ancestries^39^.

**Figure 2.**
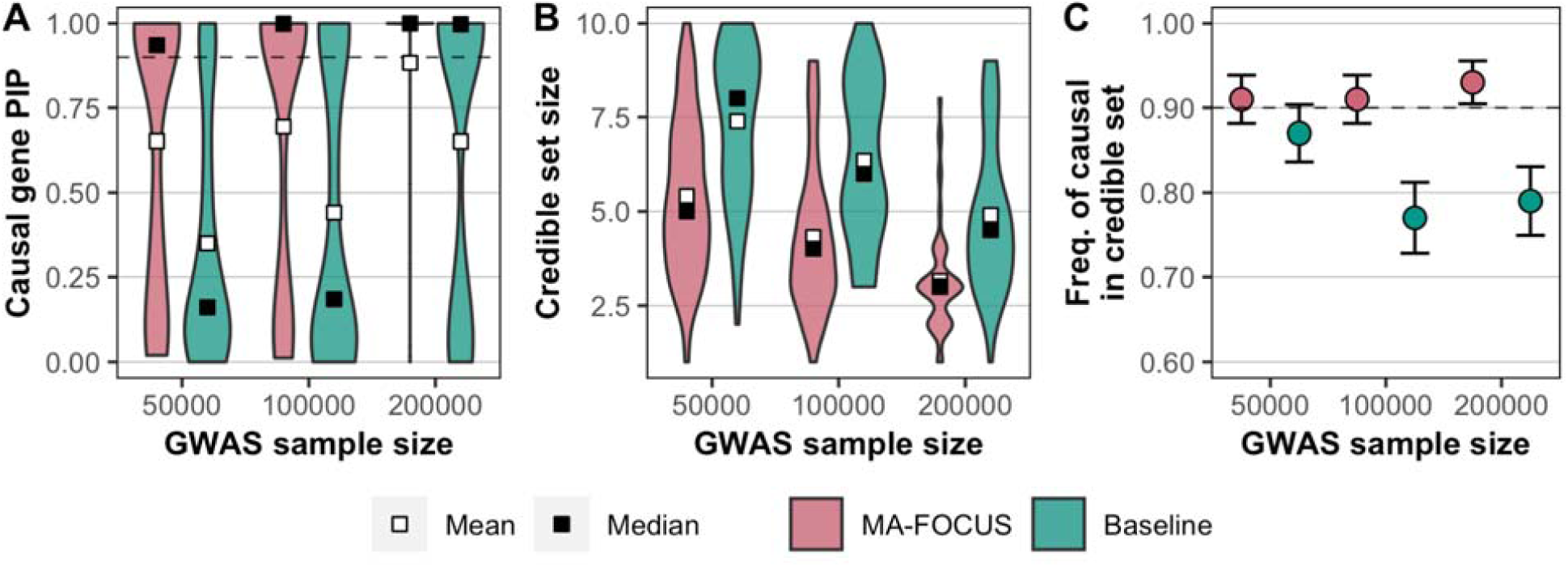
MA-FOCUS outperforms baseline approach in all three metrics as GWAS sample sizes vary when eQTLs are independent across ancestries. PIPs for 100 simulated causal genes (A), the distribution of 90% credible set sizes for 100 simulated gene regions (B), and the sensitivity (C) from MA-FOCUS, and baseline approach, varying GWAS sample sizes across multi-ancestry ancestries. See Methods section for default parameters. The black dashed lines indicate 90%. Error bars are constructed using a 95% confidence interval.

Next, we sought to quantify the increases in fine-mapping power that could be gained by including individuals from diverse genetic ancestries, rather than increasing the sample size of a single-ancestry GWAS. Specifically, we assumed an existing eQTL panel of 200 individuals for AFR and EUR ancestries and compared the performance of MA-FOCUS with single-ancestry fine-mapping, given a fixed number of total GWAS participants. We found that MA-FOCUS placed more posterior density on causal genes with a mean of 0.67 compared to 0.57 (*P* = 0.01) and produced smaller credible sets with a mean of 4.86 compared to 5.33 (*P* = 0.03) with better sensitivity of 0.91 compared to 0.83 (*P* = 0.01) when compared with FOCUS applied to equivalently powered EUR-only TWAS data (**Figure S4**). This relative performance advantage held when we compared two- to three-ancestry scenarios (**Figure S5**). Consistent with previous multi-ancestry SNP-based fine-mapping approaches^20,25^, our results suggest that incorporating additional ancestry genetic diversity in GWAS drives larger payoffs in fine-mapping performance than simply increasing the sample sizes of GWAS on previously studied ancestries.

To evaluate the performance of MA-FOCUS as a function of the underlying genetic architecture, we next performed simulations varying the *cis*-SNP heritability of gene expression 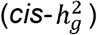 and the proportion of trait heritability attributable to a causal gene 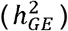. Across architectures, MA-FOCUS significantly outperformed the baseline (*P* = 2.52 × 10^−14^ for PIP metric, *P* = 7.45 × 10^−49^ for credible set metric, and *P* = 3.61 × 10^−4^ for sensitivity; **Figure S6-S7**). Moreover, when there is no causal gene effect (i.e. 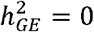), we found that MA-FOCUS returned larger PIPs for the null model (*P* = 2.88 × 10^−5^) and smaller credible sets (*P* = 1.64 × 10^−25^) on average compared with the baseline (**Figure S7**). Our results show that MA-FOCUS is better-powered than the baseline to identify the true causal model, including the null model, across a range of heritabilities for gene expression and the overall trait.

### Multi-Ancestry FOCUS is robust to genetic-architectural and data-dependent assumptions

Next, we sought to characterize the performance of MA-FOCUS when assumptions of the underlying model are partially violated. First, we simulated a complex trait where the mediating gene-trait effects differed across ancestries by setting ancestry-specific 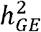 values (i.e. fixed for EUR and varying for AFR across a range; see **Methods**). Again, we found that MA-FOCUS consistently reported higher PIPs for causal genes (*P* = 3.42 × 10^−11^) and smaller 90% credible sets (*P* = 6.80 × 10^−33^) compared with the baseline (**Figure 3, S8**). Furthermore, the sensitivity of gene sets reported by MA-FOCUS were robust to up to 7-fold differences in ancestry-specific 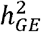 (i.e. 7.57 × 10^−4^ for EUR compared to 1.14 × 10^−4^ for AFR). Only when the AFR 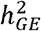 was ∼2% of the EUR 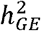 (i.e. 7.57×10^−4^ for EUR compared to 1.71 × 10^−5^ for AFR) did we find MA-FOCUS performance to degrade, which is consistent with reduced statistical power under a fixed sample size. Together these results show that MA-FOCUS is generally robust to ancestry-specific architectures.

**Figure 3.**
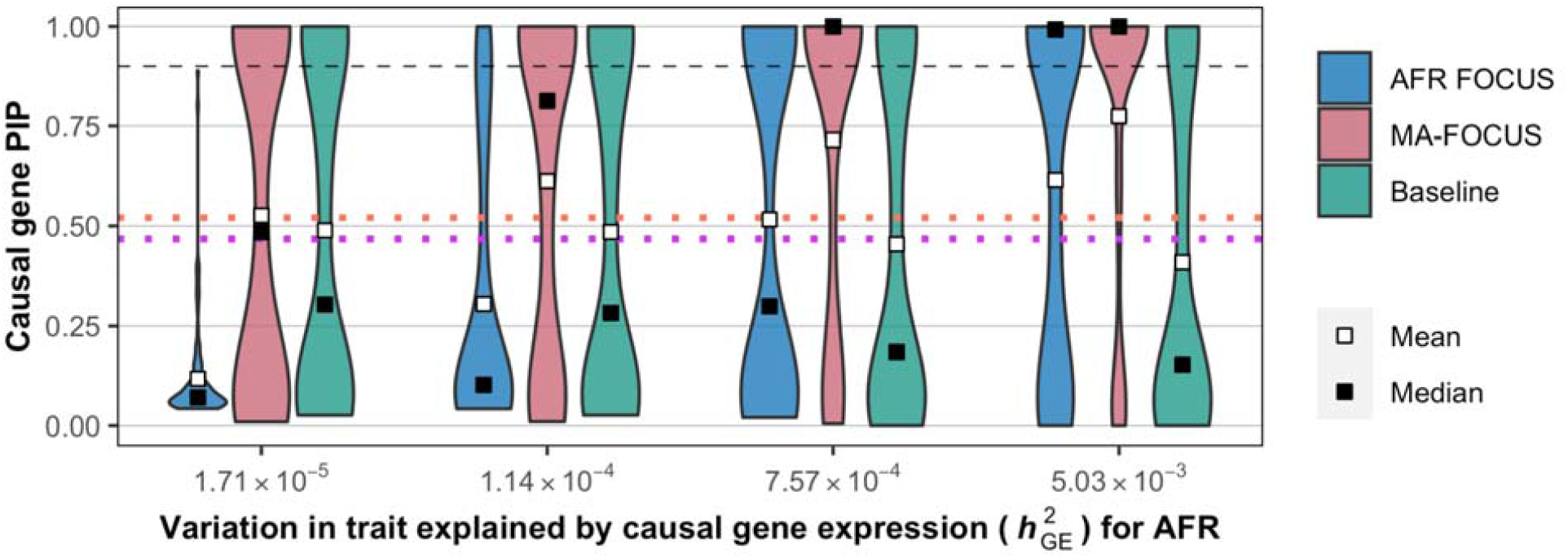
MA-FOCUS remains robust in having higher causal gene PIPs when trait heritability mediated by gene expression differs across ancestries. Distribution of inferred PIPs at the causal gene when the trait architecture varies across ancestries. We fixed trait variation explained by causal gene expression to for simulated EUR individuals while varying its amount in AFR individuals. The orange and purple dotted line indicate the mean and the median of PIPs using EUR FOCUS. The black dashed lines indicate 90%.

To investigate the impact of imbalanced GWAS sample sizes, we performed simulations matching the sample sizes of a recent multi-ancestry blood trait GWAS^18^ (*N*_*EA*_ = 511,471 and *N*_*AA*_ = 13,298; see **Methods**). In this setting MA-FOCUS computed credible sets that were smaller compared to the baseline (*P* = 3.54 × 10^−6^; **Figure S9B**) with similar mean PIPs at the causal genes (*P* = 0.13 **Figure S9A**) and sensitivity (*P* = 0.17; **Figure S9C**). This demonstrates that, even when GWAS sample sizes vary by an order of magnitude across ancestries, MA-FOCUS provides improved fine-mapping performance.

Next, we performed simulations where the trait-relevant tissue for AFR was unavailable and was substituted with eQTL data quantified in a proxy tissue with correlated genetic effects (see **Methods**). Performance of MA-FOCUS was highly dependent on the underlying correlation between proxy and causal tissue, and increased with increasing inter-tissue genetic covariance, as expected (**Figure S10**). We again observed that MA-FOCUS outperformed the baseline approach as well as single-pop FOCUS on AFR across all metrics (*P* < 1 × 10^−7^ for all PIP and credible set metrics, and *P* = 0.02 with MA-FOCUS Baseline comparison and *P* = 0.09 with MA-FOCUS AFR-FOCUS comparison for sensitivity; **Figure S10**).

Finally, we performed simulations where eQTL reference panels for AFR are not available and EUR weights are used instead for both TWAS and fine-mapping. We found that MA-FOCUS’ relative performance was mixed across different metrics, estimating similar causal PIPs and sensitivity (*P* = 0.69 and *P* = 0.90; **Figure S11A, C**) and smaller credible sets size (*P* = 3.73 × 10^−9^; **Figure S11B**). In all, this highlights the importance of multi-ancestry study design collecting gene expression data from different ancestries when possible.

### Multi-ancestry TWAS identifies shared architecture in blood traits

After confirming that MA-FOCUS outperforms other methods of TWAS fine-mapping, we next sought to apply it to real data from cohorts of European- (EA) and African-ancestries (AA) ancestries. We performed ancestry-matched TWAS for 15 blood traits using GWAS summary statistics^18^ (**Table S2, S5;** N_EA_=511,471, N_AA_=13,298) together with an eQTL reference panel of LCLs from the GENOA study^39^ (eQTL: N_EA_=373, N_AA_=441; see **Methods**). First, we estimated cis-genetic variance 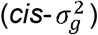 and SNP-heritability 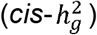 for expression at 14,797 genes assayed in both EA and AA GENOA cohorts (see **Methods**). We observed that, across all genes, 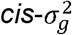 was significantly non-zero with an average of 0.018 for EA compared to 0.024 for AA *(P* < 1 × 10^−100^ for both tests). Furthermore, focusing on the 4,646 genes whose expression was significantly heritable in at least one of the cohorts, 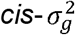 estimates were positively correlated across ancestries with *r* = 0.54 (*P* < 1 × 10^−100^ for both tests against 0 and 1; **Figure 4A**), which is consistent with previous results suggesting that the genetic architecture of gene expression is significantly shared across ancestries^39^. Next, we trained prediction models using the FUSION pipeline and performed 5-fold cross-validation (CV; see **Methods**). We found CV *r*^2^ was significantly non-zero (AA CV *r*^2^ =0.11; EA CV *r*^2^=0.10; < 1 × 10^−100^ for both), which were strongly correlated with 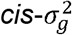 estimates (r=0.72 for EA with P < 1 × 10^−100^; *r*=0.76 for AA with P < 1 × 10^−100^; **Figure S12AB**), suggesting that in-sample prediction models perform well and are consistent with theory where heritability provides a predictive upper bound^39,58,59^.

**Figure 4.**
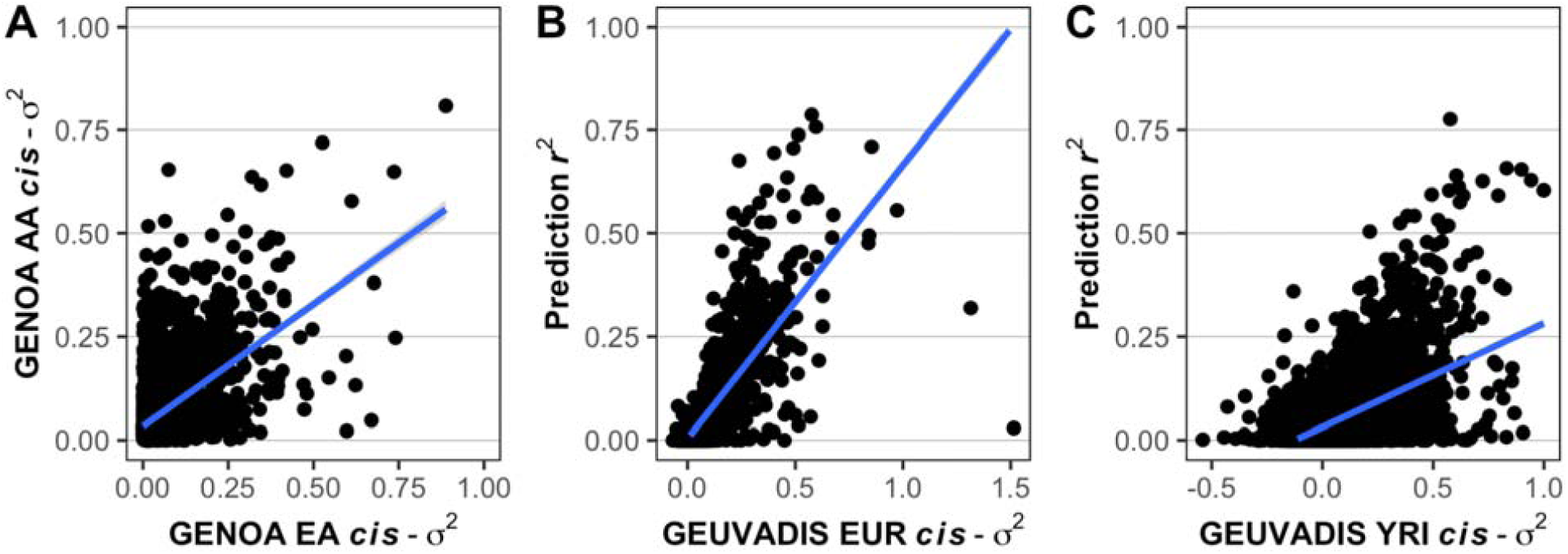
Heritability and correlation analysis reveal evidence for shared genetic architecture for expression in LCLs. (A). The scatter plot for the genetic variance (*cis-*) of LCL gene expression for AA and EA ancestry in GENOA study. (B) (C). The scatter plots where the y-axis is a squared correlation between measured LCL gene expression in GEUVADIS and predicted by eQTL panels from GENOA, and x-axis is *cis-*. Each point represents a gene. The blue line is estimated using ordinary linear regression.

Next, we further validated the predictive performance of LCL expression models by evaluating their out-of-sample performance in the European- and Yoruba-ancestry cohorts (EUR and YRI, compared with GENOA EA and GENOA AA, respectively) of the independent GEUVADIS study (see **Methods**)^53^. While YRI are not an ideal ancestry proxy for admixed African Americans, we expect a significant degree of genetic similarity between the two given the high mean West-African component of African Americans (∼80%), which YRI is commonly used to represent^39^. Consistent with our GENOA-based findings, we found that the estimates of 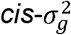 in GEUVADIS were significantly non-zero with an average of 0.062 for EUR compared to 0.077 for YRI (*P* < 1 × 10^−100^ for both tests). We then calculated the *r*^2^ between measured LCL gene expression from GEUVADIS individuals and expression predicted using our GENOA-based weights. We found an average out-of-sample *r*^2^ estimate of 0.04 for EUR and 0.05 for YRI (*P* < 1 × 10^−100^ for both tests**; Figure S13**), which while decreased compared to within-GENOA estimates, suggests our predictive models accurately capture the genetic component of gene expression for ancestry groups. Estimates were significantly correlated with estimates of GEUVADIS 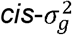, with *r* = 0.79,0.54 for EUR and YRI (*P* < 1 × 10^−100^ for both tests against 0; **Figure 4BC**). The comparatively poorer 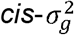-adjusted performance of our AA expression weights in the GEUVADIS YRI is not unexpected, given the ancestry differences between the Yoruba and African-Americans, discussed above, which likely impact the genetic regulation of gene expression. Indeed, we found that 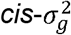 for AA and YRI are less correlated than EA and EUR (*r*=0.30 and 0.55 with *P* < 1 × 10^−100^ for both tests against 0; *P* < 1 × 10^−40^ for testing correlation difference)^60^. Next, we evaluated across-ancestry prediction performance by predicting LCL gene expression levels for GEUVADIS EUR individuals using GENOA AA weights (similarly for GEUVADIS YRI and GENOA EA) and estimated an average *r*^2^ 0.039 for EUR and 0.033 for YRI (*P* < 1 × 10^−100^ for both tests; **Figure S13**). Consistent with previous works^59^, we found a decrease in accuracy for GEUVADIS YRI individuals compared to within-ancestry results (*P* = 1.75 × 10^−31^) and similar levels of accuracy for GEUVADIS EUR (*P* = 0.09). Together, these results demonstrate that our prediction models capture accurately the heritable component of gene expression within ancestry groups and recapitulate previous findings on the limited transportability of cross-ancestry prediction models for gene expression^39,59^.

Having validated our SNP-based LCL expression prediction models, we next conducted multi-ancestry TWAS for each of the 15 blood traits on 4,579 heritable genes in 995 unique independent regions (see **Methods**). Across all traits, we identified a total of 6,236 (2,009 unique), 116 (57 unique) genome-wide TWAS significant genes in EA and AA, respectively, of which 28 were shared (17 unique) in 3029 (623 unique) regions (*P* < 0.05/4579; 3.29 unique genes per region; **Figure 5A**; **Table S6;** see **Data Availability** for the full results). Of the 8,416 (1064 unique) LD blocks that contain GWAS signal (*P* <5 × 10^−8^) in either ancestry or the meta-analysis, 2,933 (623 unique) also exhibited TWAS signals. Conversely, 96 (78 unique) LD blocks that contain TWAS signal do not exhibit GWAS signals, suggesting that TWAS identified novel risk regions for 15 blood traits. Of the 3,029 (623 unique) regions containing TWAS hits, 1,335 (319 unique) contain multiple TWAS significant associations, motivating the use of gene fine-mapping. We found that both normalized GWAS and TWAS effect size correlations between EA and AA are significantly non-zero for all traits, suggesting shared architecture at the individual SNP- and gene-effect level (**Table S7**; **Figure S14;** see **Methods**). Interestingly, we found that across-ancestry correlations are 20% higher on average for TWAS compared to GWAS (*r* = 0.061 and 0.052, respectively, *P* = 0.027; **Figure 5B**; **Table S7**), which is consistent with previous findings demonstrating that predicted transcriptomic risk scores better correlate across ancestry groups^63^ and suggests that gene-level effects on average better reflect shared biology compared with SNP-level effects.

**Figure 5.**
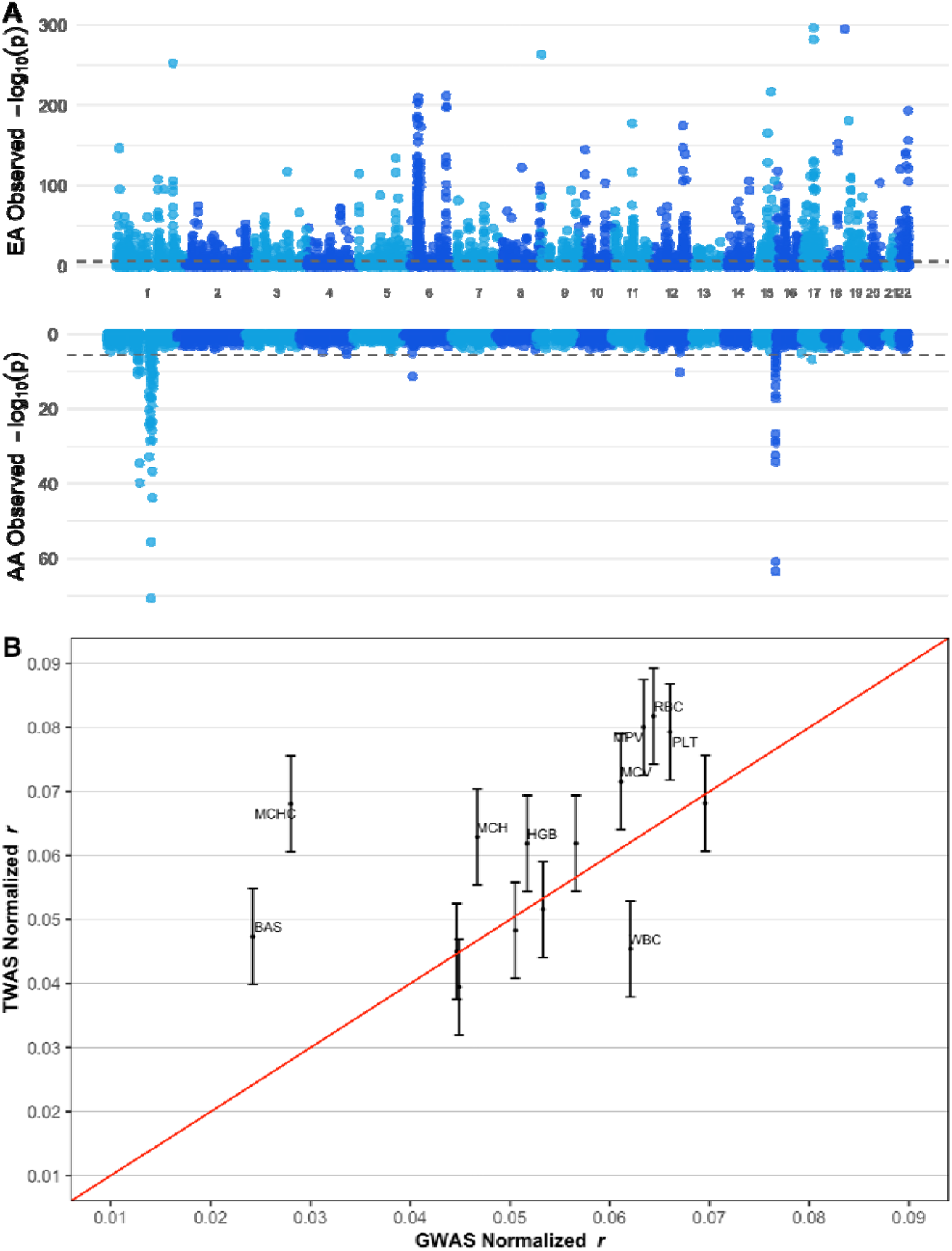
The TWAS Manhattan Plot indicates highly correlated genes at certain regions. (A). The upper plot is the Manhattan plot for EA TWAS, and the lower is for AA TWAS across all 15 traits. Colors differentiate adjacent chromosomes. (B). Cross-ancestry correlation of TWAS and GWAS effect sizes (see **Methods**). The correlations are higher on average for TWAS compared to GWAS (r = 0.061 and 0.052, respectively,). Each point represents a trait and the red line is the identity line.

### Trans-ancestry fine-mapping prioritizes likely causal genes in blood traits

Next, we applied MA-FOCUS to TWAS results for 10 blood traits focusing on 163 genes overlapping the 11 unique regions that contained TWAS signals for both EA and AA ancestry for a given trait (see **Methods**). Across these 11 regions, each contained on average 7.45 TWAS significant associations and 3.05 genes in the credible set, none of which contained the null-model. We estimated an average 2.85 causal genes per region by summing over local PIPs in credible sets, with 20 out of 22 credible sets containing three or fewer genes (**Table S8;** see **Data Availability**). The average maximum PIP across credible sets was 0.99 (SD=0.03) and retained similar PIPs for second, and third rank (**Figure S15**). While estimated PIPs across methods were correlated, when comparing the credible sets output by MA-FOCUS and other multi-ancestry approaches, we observed higher means and smaller standard deviations of PIPs for MA-FOCUS than other approaches (**Figure 6, S16**). For 67 trait-gene pairs in MA-FOCUS credible gene sets, 59 are not detected by EA-FOCUS; out of 22 top genes in credible sets, 7 are not detected by EA-FOCUS, respectively, which suggests that incorporating non-European data in well-powered loci can prioritize additional putative causal genes (**Figure S17**). Next, to determine the extent to which prioritized genes are likely to be shared or ancestry-specific, we performed model comparison using Bayes factors from MA-FOCUS and FOCUS (see **Methods**). We observed an average log-scale BF of 1.48, suggesting that credible-set genes underlying these blood traits are much more likely to be shared across ancestries than ancestry-specific (**Figure S18**).

**Figure 6.**
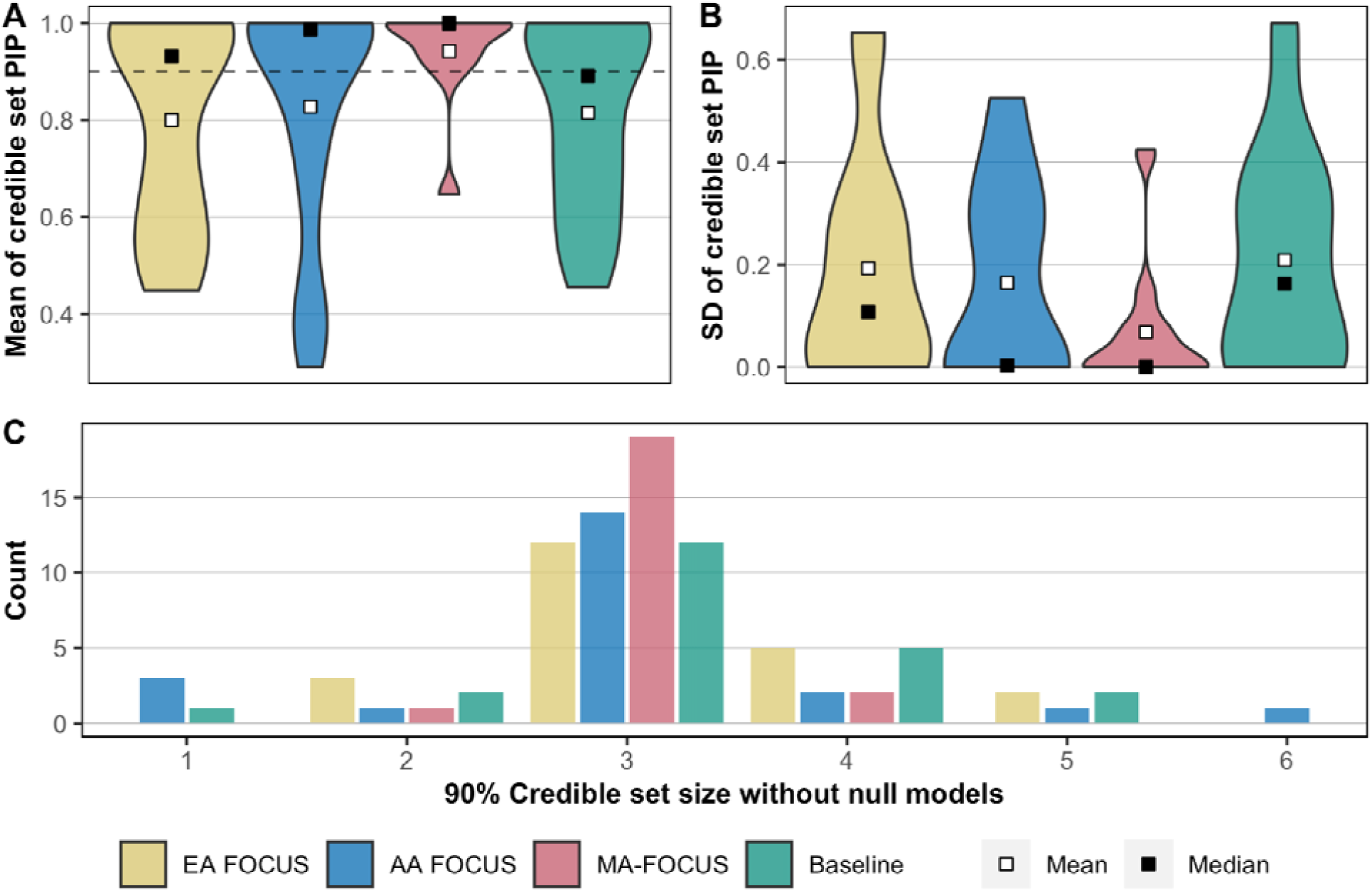
Credible sets output by MA-FOCUS have higher mean PIPs and lower standard deviation while exhibiting similar credible set size of EUR FOCUS and the baseline approach. The distribution of (A) the mean of gene PIPs in credible sets, (B) the standard deviation (SD) of gene PIPs in credible sets, (C) credible set size. Calculations do not include null models. Mean and median are represented by blank and black boxes for the metric distribution.

Next, we investigated genes to which MA-FOCUS assigned a high PIP (> 0.75) and included in a credible set, but that were not identified by the baseline approach. We refer to these genes hereafter as the ‘MA-FOCUS-specific genes’. We also looked at the converse situation: that is, genes that the baseline approach found strong support for, but that were not prioritized by MA-FOCUS, and refer to these at the ‘baseline-specific genes’. Importantly, we found that all 21 baseline-specific genes had low PIPs (< 0.1) from ancestry-specific fine-mapping in at least one ancestry, while 10 of these genes had a low PIP in both ancestries. On the other hand, only two out of 31 total MA-FOCUS-specific genes had PIPs below 0.1 in both AA and EA. Five out of 31 total MA-FOCUS-specific genes achieved a moderate PIP of at least 0.4 in both EA and AA ancestry-specific fine-mapping (**Figure S19**). These five genes are *ARNT, BAK1, NPRL3, PHTF1*, and *TARS2*. A literature search uncovered additional evidence for roles in cardiovascular system disease and development (specifically, blood cell and vasculature formation, diabetes, leukemia, and cardiomyopathy) among these MA-FOCUS-specific genes (**Figure S20**)^61,62,64-69,71,72^. Overall, this result suggests that by appropriately modelling across-ancestry heterogeneity, MA-FOCUS can prioritize disease-relevant genes that would otherwise be missed from naïve meta-analyses.

To validate genes prioritized by MA-FOCUS and the baseline approach, we next performed a series of validation tests comparing the credible sets (see **Methods**). First, we performed gene set enrichment analysis on the credible set genes using the DisGeNET dataset across all 15 blood traits. We found that MA-FOCUS’ credible sets are enriched more in hematological measurement categories compared to the baseline approach (25 and 13 categories, meta-analysis P-value of 1.73 × 10^−16^ compared to 2.91 xlO^−11^; **Figure 7**; **Table S9**). Second, by restricting our focus to trait-matched DisGeNET enrichment categories, we observed that MA-FOCUS output more significantly enriched credible gene sets compared to the baseline approach (meta-analysis P-value of 7.56 × 10^−5^ compared to 1.3 × 10^−3^; **Figure 7**; **Table S10**). Third, using curated “silver standard” databases consisting of Online Mendelian Inheritance in Man (OMIM) and Orphanet for 159 blood-related diseases (see **Web Resources**; see **Methods**), we observed MA-FOCUS outputs a higher average AUROC curve with 0.57 compared to 0.43, suggesting improved performance in predicting causal genes of monogenic and oligogenic blood-related Mendelian and rare diseases (**Table S11**). Altogether, we find that credible set genes computed using MA-FOCUS better reflect relevant disease biology compared to single-ancestry and alternative approaches.

**Figure 7.**
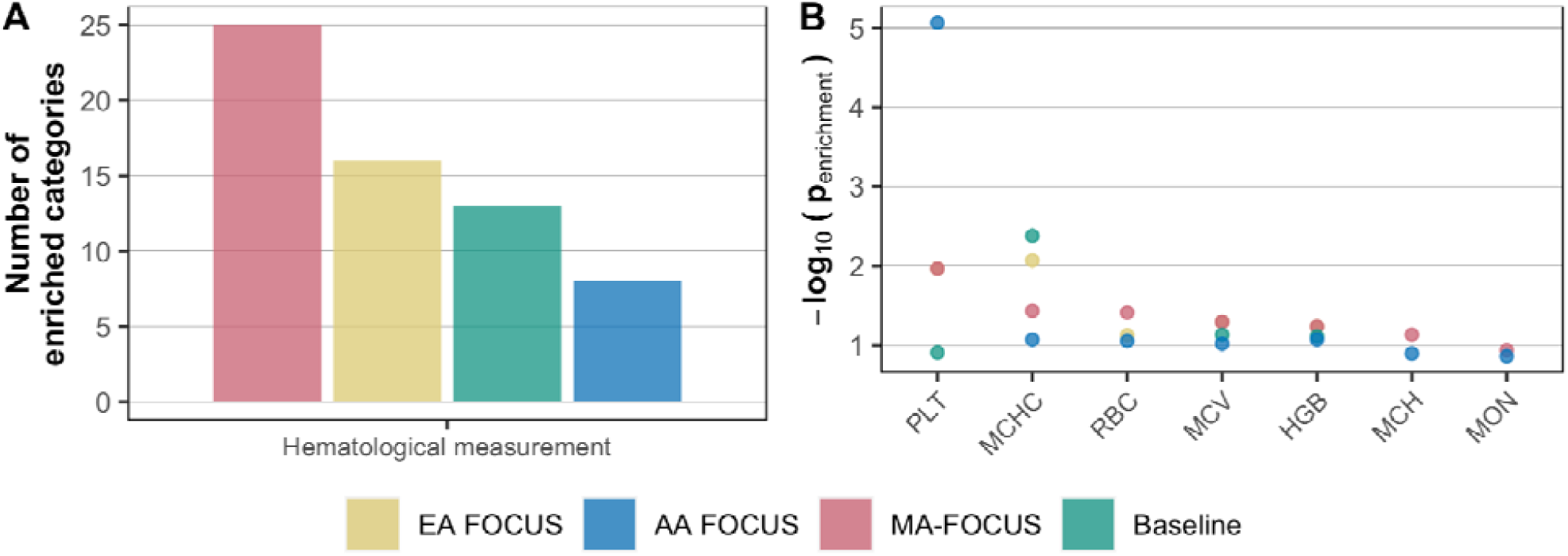
Genes prioritized by MA-FOCUS are enriched in hematological categories more often than other methods. (A) The bar plot shows the number of enriched categories in DisGeNET identified by each method within the hematological-measurement-related category. The enriched category is defined as Bonferroni-corrected P-value less than 0.05. (B) The dot plot shows enrichment by categories in DisGeNET corresponding to 7 blood traits.

## Discussion

In this work, we present MA-FOCUS, a Bayesian fine-mapping method that incorporates GWAS and eQTL data together with LD reference panels from multiple ancestries of diverse genetic ancestries to estimate credible sets of causal genes for complex traits. Our method is unique in that it explicitly accounts for, and takes advantage of, heterogeneity in LD and the genetic architecture of gene expression to improve TWAS fine-mapping performance. Importantly, our method assumes only that causal genes for complex traits are shared across ancestries while making no assumptions on underlying eQTL architectures across ancestries. This is an important feature of our method considering recent findings that SNP-level replication across genetic ancestries is weaker than gene-level replication^36^, and that only ∼30% of SNP-gene expression associations are shared between European- and African-American ancestry^39^. Through extensive simulations, we demonstrate that MA-FOCUS’ ability to identify causal genes is superior to baseline approaches and is robust to data-dependent limitations (see **Methods**).

We perform ancestry-specific TWAS and apply MA-FOCUS to 15 blood traits using GWAS statistics in Chen et al. and lymphoblastoid cell line eQTL data in GENOA from cohorts of primarily European and African continental ancestry. We report 3.29 TWAS significant genes per region in 623 regions across all blood traits. The cross-ancestry heritability analysis on LCL gene expression data, together with correlation analysis on blood traits of GWAS and TWAS statistics, recapitulate evidence for shared genetic architecture of blood traits between the two ancestries, and provide evidence for gene-level effects correlating better across ancestries compared with SNP-level effects. Next, in 22 regions that contain TWAS signals for both ancestries, MA-FOCUS reports 3.05 genes in the credible sets and estimated 2.85 putative causal genes per region across all blood traits. We validate MA-FOCUS’ credible sets by performing enrichment analyses and referencing the results of functional studies. We show that MA-FOCUS’ credible sets are more strongly enriched for relevant genes associated with hematological traits in the DisGeNET platform, a database of genotype-phenotype associations compiled from various sources (**Figures 7**). Importantly, MA-FOCUS identifies genes that are known to have functional relevance for cardiovascular system disease and development but are not identified by the baseline approach.

Despite MA-FOCUS’ advantages in performance, as demonstrated through extensive simulations, we note several limitations to our analysis of blood traits. First, we find that MA-FOCUS’ performance advantage is attenuated when the EA sample size is approximately 40 times greater than the AA sample size (**Figure S6**). Across the 10 blood traits evaluated for fine-mapping, all methods outputted similarly sized 90% credible sets (**Figure 6**). Additionally, the MA-FOCUS’ PIPs are overall strongly correlated with all three of these approaches (**Figure S16**). Despite this, as discussed previously, we find evidence that MA-FOCUS is more successful than other approaches at identifying genes that are functionally associated with blood traits. Secondly, the gene expression data for our eQTL reference panel was derived from immortalized cell lines^39^ which differ from complex living organisms in fundamental ways. Therefore, this tissue type may not be the most appropriate tissue for identifying causal relationships with blood traits. When we explored this scenario using simulations, we found that both causal gene PIPs and calibration were substantially reduced when poorly correlated tissue was used for one of the ancestries (**Figure S10**). We expect this effect would be exacerbated if a poorly correlated tissue was used to estimate weights for both ancestries. Thirdly, our eQTL reference panel and GWAS cohort for the AA ancestry represent genetically admixed individuals whose genomes are a combination of (West) African and European ancestry. Therefore, when estimating weights for this ancestry, the local ancestry at any given locus would include some proportion of European-derived genotypes. This likely introduces noise and further reduces the power of our weight estimates. In total, our analysis limitations motivate us to perform large-scale GWAS and eQTL study on non-European ancestry and admixed populations with comprehensive types of tissues and cell-types.

Here, we describe general caveats of our multi-ancestry TWAS fine-mapping approach. First, MA-FOCUS assumes that genes causal for complex traits are shared across ancestries, which neglects the possibility of ancestry-specific causal genes. However, because several large-scale multi-ancestry GWAS studies have shown that the majority of risk signals replicate in ancestries, we believe this to be a relatively minor issue^18-22^. Second, MA-FOCUS models complex traits as a linear combination of steady-state gene expression, which neglects potential gene-environment interaction (GxE) or gene-gene interaction (GxG). While several works have supported linear assumptions for complex traits through large-scale GWAS results^58,70^, recent work analyzing large-scale GWAS from multiple ancestries has provided evidence that allelic heterogeneity across ancestries may be due to GxE^19^, and we acknowledge this as a potential interesting direction.

Overall, MA-FOCUS provides Bayesian inference on gene causality for complex traits in specific genomic regions leveraging GWAS data, eQTL data, and LD data of multiple ancestries. It improves the precision in gene fine-mapping by accounting eQTL and LD heterogeneity across different ancestral groups and sheds light on the genetic architecture of complex traits.

## Appendix

### Estimating TWAS causal effect prior variance

Here we describe an estimator for the prior causal effect-size variance (i.e. 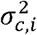) similar to the HESS model for local heritability^41^. Our model assumes that marginal TWAS *z* statistics for *m* genes have a sampling distribution given by

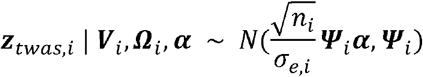

where 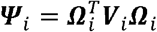. We would like to define an unbiased estimator for the variance explained by (fixed) causal effects ***α***. Specifically, *Var(G*_*i*_*α) = α*^*T*^*Var(G*_*i*_*) α = α*^*T*^*W*_*i*_^*T*^ *V*_*i*_*Wα = α*^*T*^*Ψ*_*i*_*α*. As a result, an intuitive (but biased) estimator for 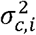 would be

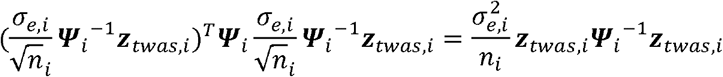

In practice 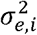 is extremely close to 1, hence an unbiased estimator for the sample-size scaled causal effect prior variance 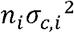 is given by

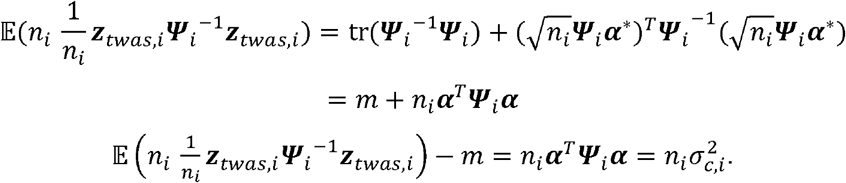

In practice when the estimator is negative (e.g., when little TWAS signal exists), we use the biased estimator to ensure positivity.

## Acknowledgements

We are thankful to Drs. Sharon Kardia and Jennifer Smith for providing resources to enable model fitting in GENOA eQTL data. GENOA genotype and gene expression data were supported by grants from NHLBI (HL054457, HL054464, HL054481, HL119443, and HL087660). This work was funded in part by National Institutes of Health (NIH) under awards R01HG012133 and R01GM140287.

## Web resources

PLINK: https://www.cog-genomics.org/plink/

MESH: https://www.nlm.nih.gov/mesh/meshhome.html

Bedtools: https://bedtools.readthedocs.io/en/latest/

FUSION: http://gusevlab.org/projects/fusion/

GCTA: https://cnsgenomics.com/software/gcta/

Silver standard TWAS analysis: https://github.com/hakyimlab/silver-standard-performance

UpsetR: https://github.com/hms-dbmi/UpSetR

EnrichR: https://cran.r-project.org/web/packages/enrichR/index.html

## Data and code availability

MA-FOCUS software: https://github.com/mancusolab/focus

LCL prediction models and complete TWAS fine-mapping results: https://www.mancusolab.com/ma-focus

Analysis codes: https://github.com/mancusolab/MA-FOCUS-data-code

