## Supplemental Figures for "Multi-ancestry fine-mapping improves precision to identify causal genes in transcriptome-wide association studies"

* contributed equally

** corresponding author

### **Supplemental Figures**

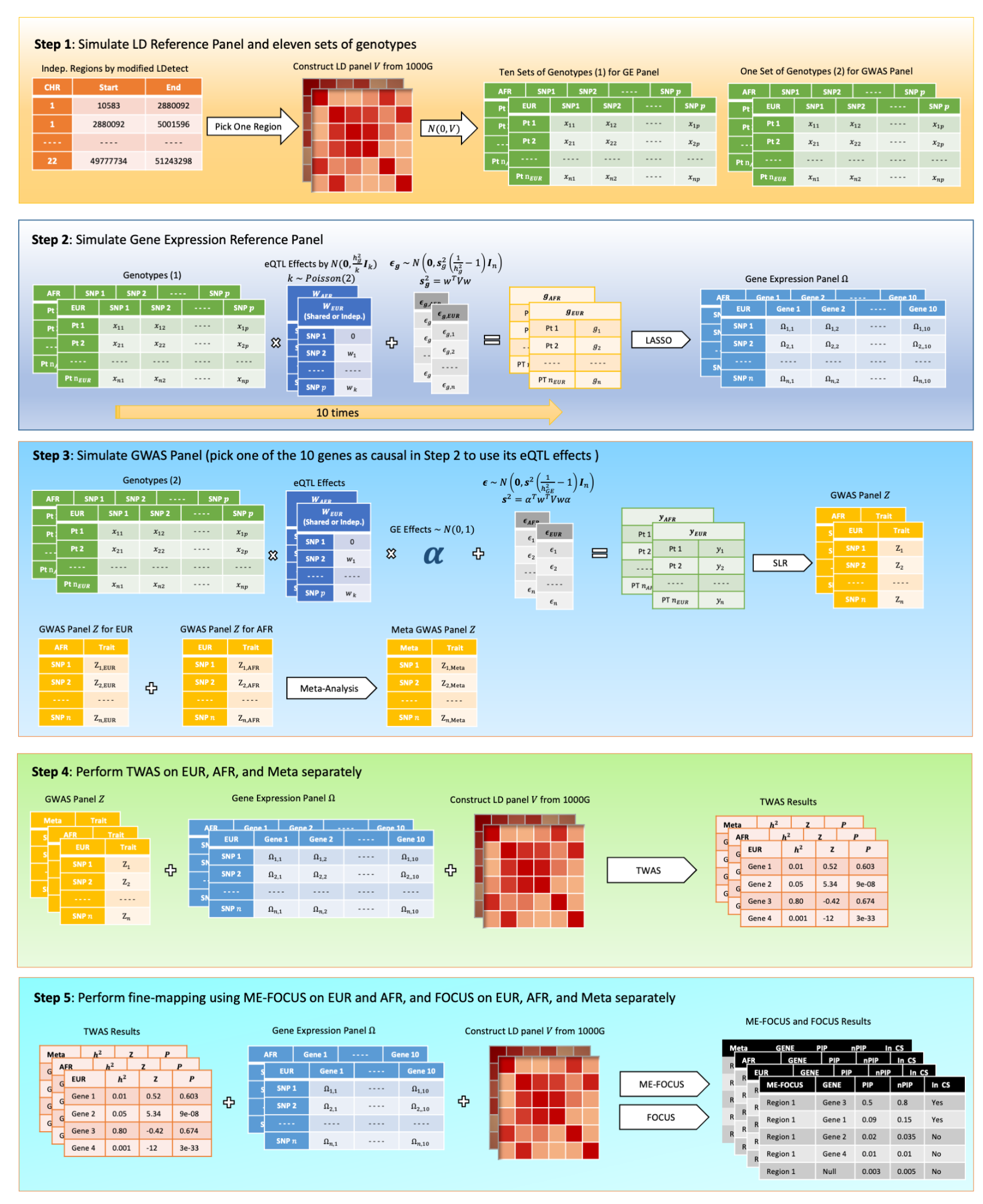

##### **Figure S1. Simulation Process Illustration.**

General procedure of the simulation consists of 5 steps. First, we computed approximately independent LD blocks for two ancestries. Second, we constructed the eQTL reference panels by sampling genotypes from the LD blocks, simulating eQTL effects, and computing gene expression at causal and non-causal genes. Third, we calculated the GWAS summary statistics by sampling genotypes and simulating a complex trait as a function of eQTL effects of the causal gene from the second step. Fourth, we performed a TWAS using penalized models fitted in the eQTL reference panels. Fifth, we performed fine-mapping using different approaches, including our new method, MA-FOCUS.

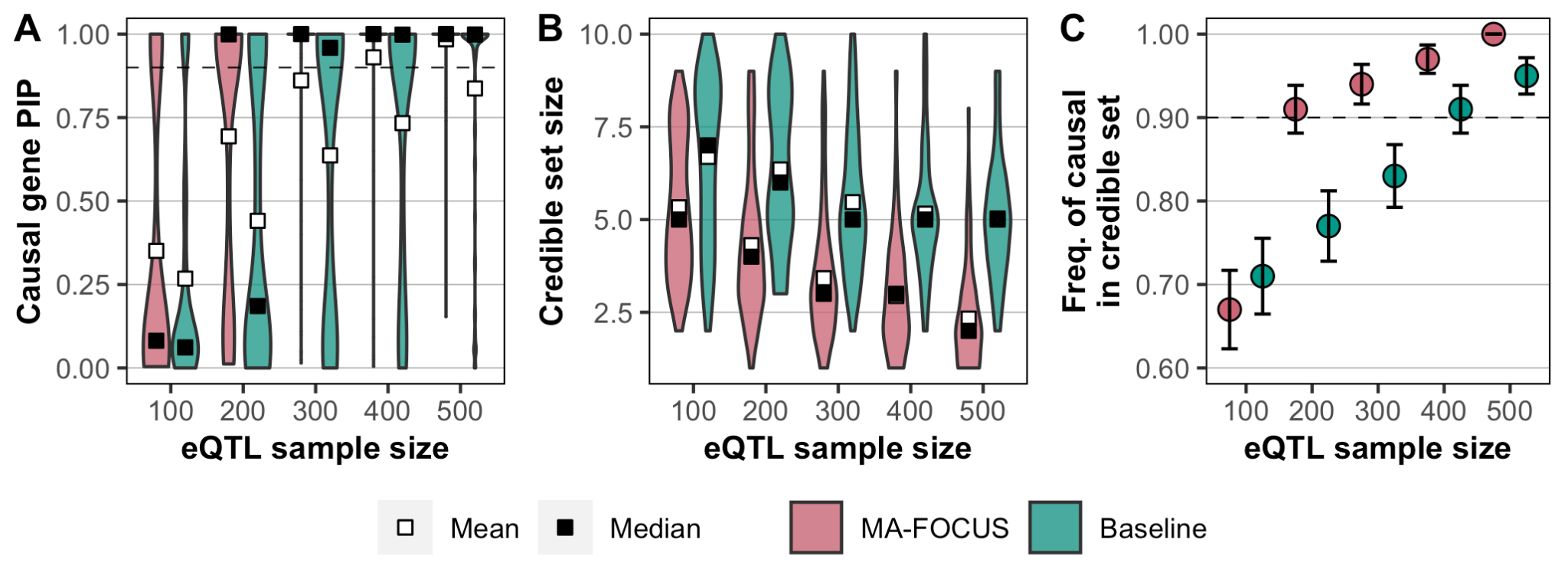

##### **Figure S2. MA-FOCUS outperforms the baseline approach in all three metrics as eQTL sample sizes vary when eQTLs are independent across ancestries.**

PIPs for 100 simulated causal genes (A), the distribution of 90% credible gene set sizes for 100 simulated gene regions (B), and the sensitivity (C) from MA-FOCUS and the baseline approach, varying eQTL panel size for both ancestries. See Methods section for default parameters. Error bars are constructed using a 95% confidence interval.

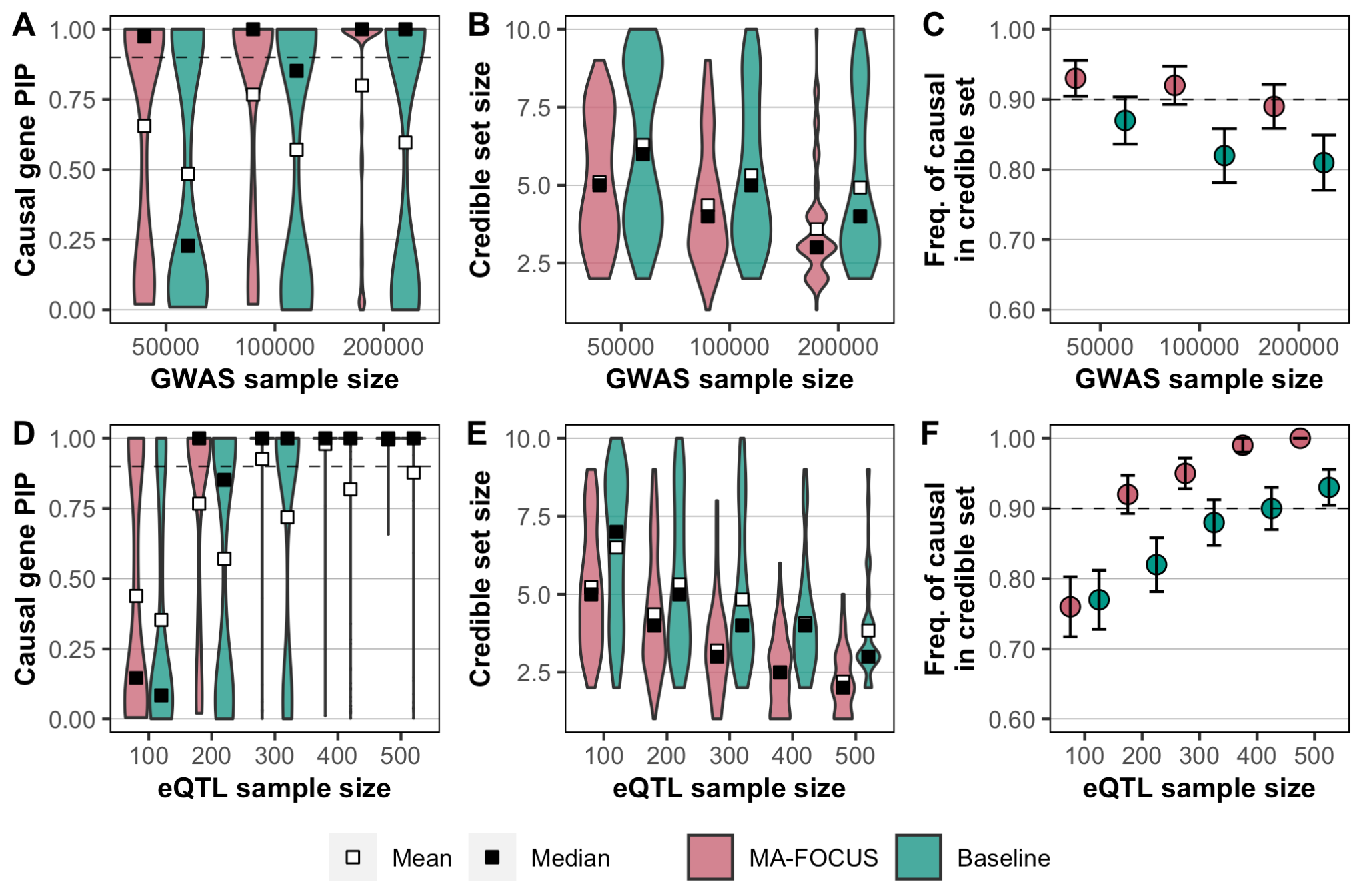

##### **Figure S3. MA-FOCUS outperforms the baseline approach in all three metrics as GWAS and eQTL sample sizes vary when eQTLs are shared across ancestries.**

PIPs for 100 simulated causal genes (A, D), the distribution of 90% credible gene set sizes for 100 simulated gene regions (B, E), and the sensitivity (C, F) from MA-FOCUS, and baseline approach, varying GWAS (A, B, C) and eQTL (D, E, F) sample sizes. The GWAS sample size was fixed at 100,000 when varying eQTL sample size, and the eQTL sample size was fixed at 200 when varying GWAS sample size. See Methods section for default parameters. The black dashed lines indicate 90%. Error bars are constructed using a 95% confidence interval.

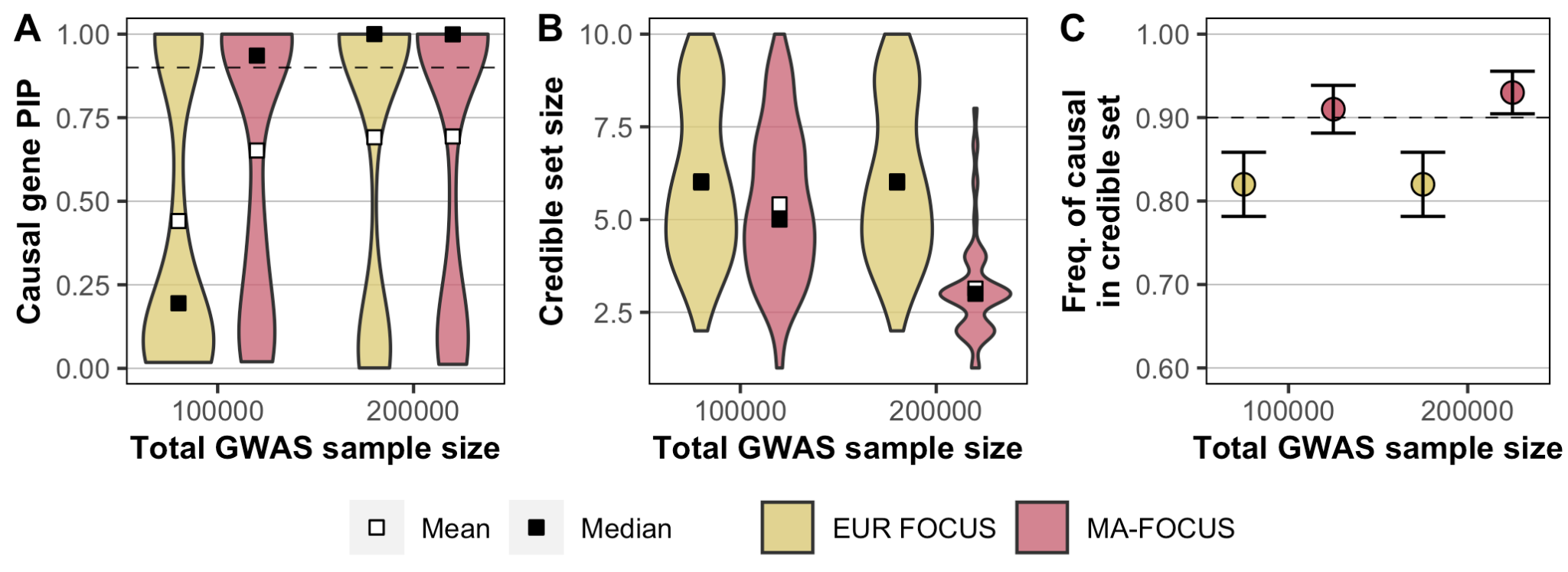

##### **Figure S4. MA-FOCUS outperforms single-ancestry FOCUS in all three metrics when total GWAS samples are fixed and eQTLs are independent across ancestries.**

PIPs for 100 simulated causal genes (A), the distribution of 90% credible gene set sizes for 100 simulated gene regions (B), and the sensitivity (C) from MA-FOCUS and the single-ancestry FOCUS for EUR, varying total GWAS sample sizes. For the MA-FOCUS approach, total GWAS sample size is equally split for each ancestry. See Methods section for default parameters. The black dashed lines indicate 90%. Error bars are constructed using a 95% confidence interval.

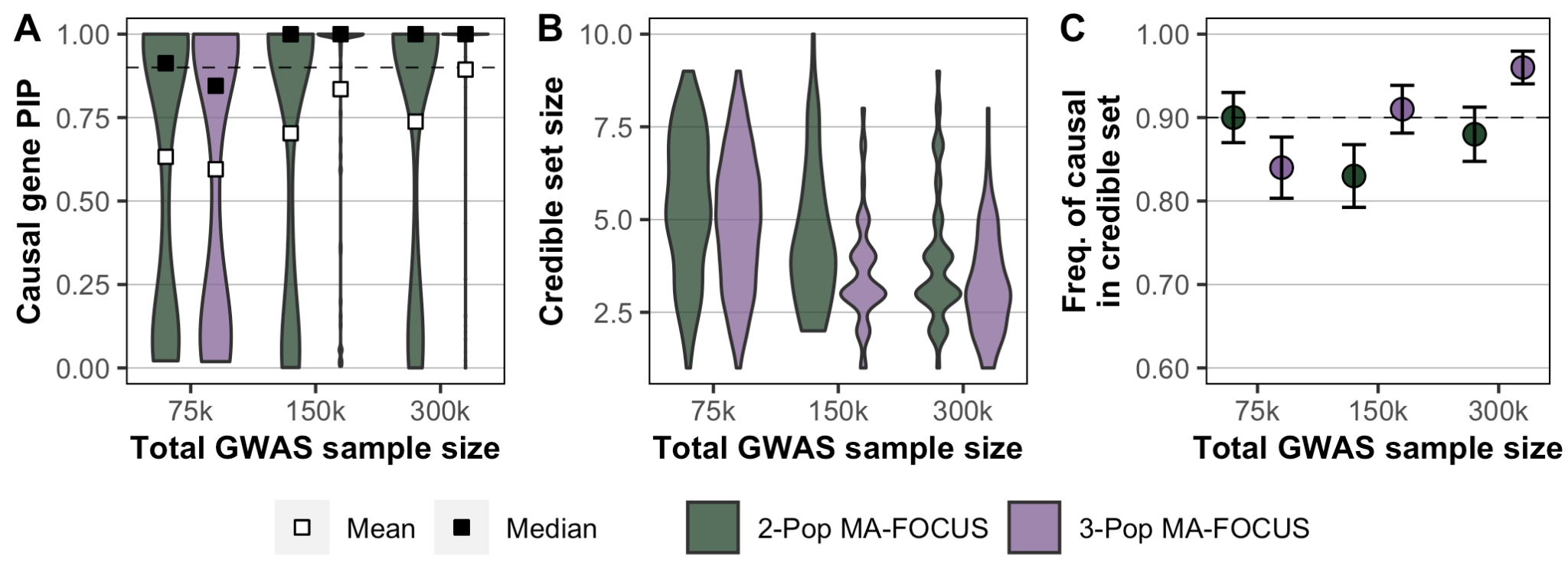

##### **Figure S5. MA-FOCUS performs better using three ancestries compared to two when eQTLs are independent across ancestries.**

PIPs for 100 simulated causal genes (A), the distribution of 90% credible gene set sizes for 100 simulated gene regions (B), and the sensitivity (C) from EUR-AFR (two-ancestry) MA-FOCUS and EUR-AFR-EAS (three-ancestry) MA-FOCUS, varying total GWAS sample sizes. For each MA-FOCUS approach, total GWAS sample size is equally split for each ancestry. See Methods section for default parameters. The black dashed lines indicate 90%. Error bars are constructed using a 95% confidence interval.

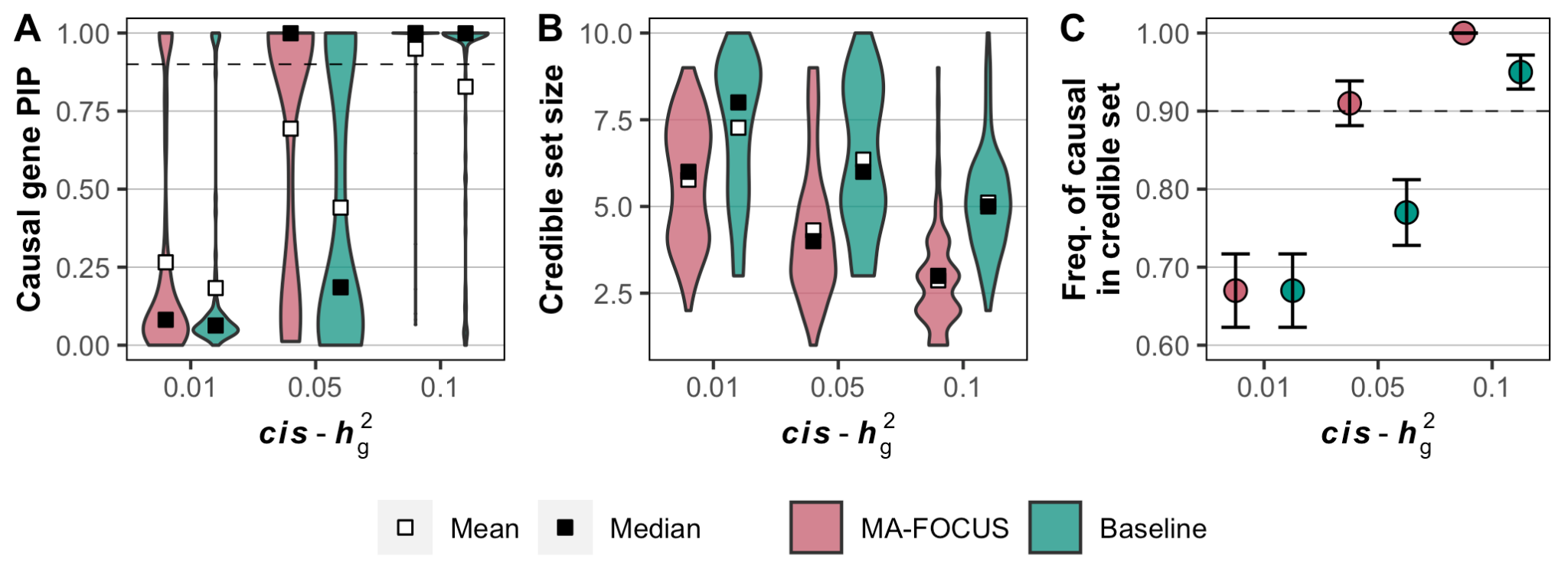

##### **Figure S6. MA-FOCUS outperforms the baseline approach while varying *cis*-**$\boldsymbol{h}_{\boldsymbol{g}}^{\boldsymbol{2}}$ **when eQTLs are independent across ancestries.**

PIPs for 100 simulated causal genes (A), the distribution of 90% credible gene set sizes for 100 simulated gene regions (B), and the sensitivity (C) from MA-FOCUS and the baseline approach, varying *cis*-$h_{g}^{2}$ for both ancestries. See Methods section for default parameters. The black dashed lines indicate 90%. Error bars are constructed using a 95% confidence interval.

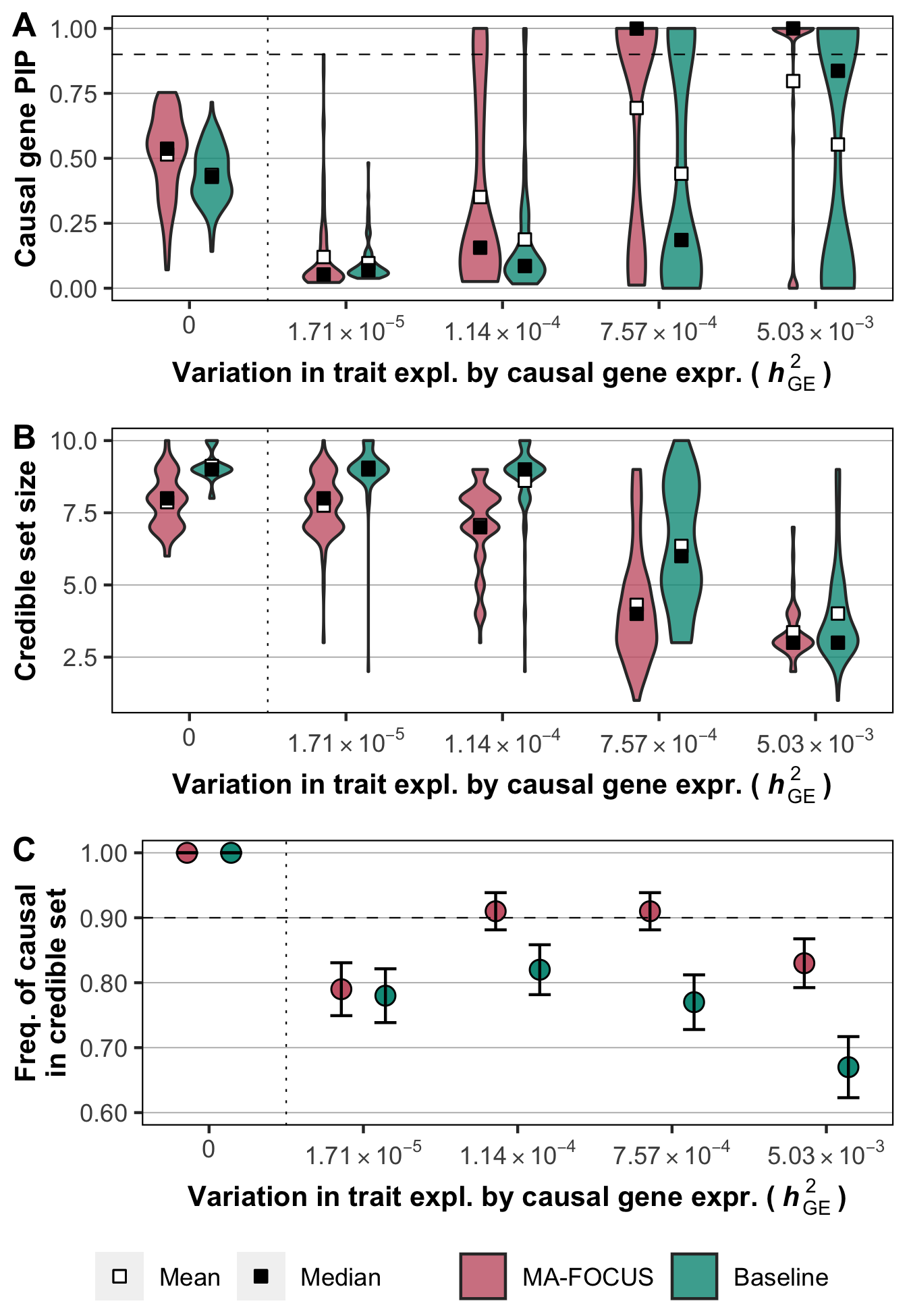

##### **Figure S7. MA-FOCUS outperforms the baseline approach while varying** $\boldsymbol{h}_{\boldsymbol{GE}}^{\boldsymbol{2}}$ **when eQTLs are independent across ancestries.**

PIPs for 100 simulated causal genes (A), the distribution of 90% credible gene set sizes for 100 simulated gene regions (B), and the sensitivity (C) from MA-FOCUS and the baseline approach, varying $h_{GE}^{2}$ for both ancestries. See Methods section for default parameters. The black dashed lines indicate 90%. Error bars are constructed using a 95% confidence interval.

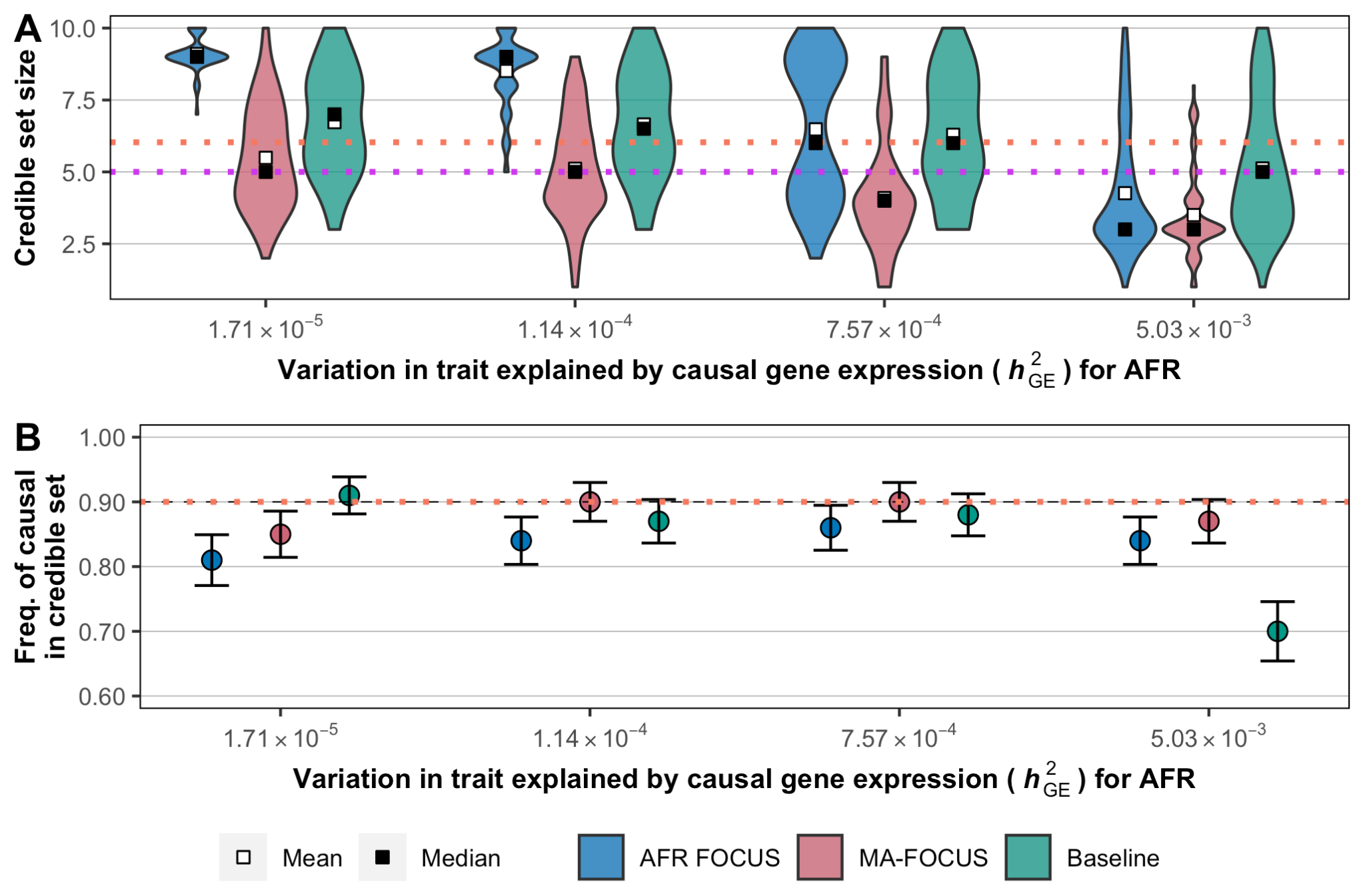

##### **Figure S8. MA-FOCUS performs robustly when** $\boldsymbol{h}_{\boldsymbol{GE}}^{\boldsymbol{2}}$ **differs across ancestries.**

The distribution of 90% credible gene set sizes for 100 simulated gene regions (A), and the sensitivity (B) from AFR FOCUS, MA-FOCUS, and baseline approach, varying complex trait variation explained by causal gene expression ($h_{GE}^{2}$) for AFR and fixing for EUR whose $h_{GE}^{2}=7.57\times10^{-4}$. See Methods section for default parameters. The orange and purple dotted line in (A) indicate the mean and the median of credible gene set size using EUR FOCUS. The orange dotted line in (B) indicates the sensitivity using EUR FOCUS. Error bars are constructed using a 95% confidence interval.

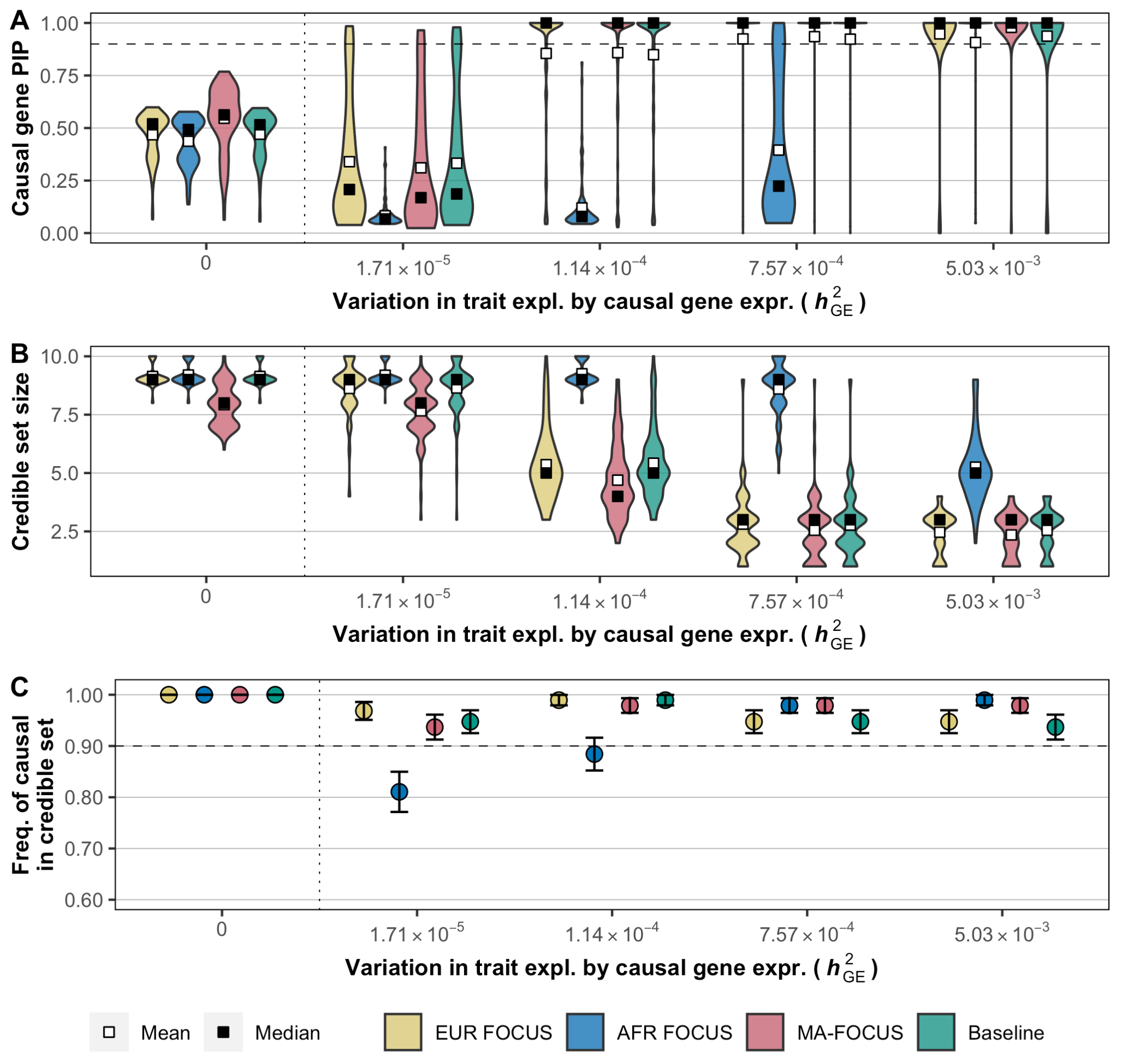

##### **Figure S9. MA-FOCUS outputs smaller gene set sizes when there exists a huge imbalance in GWAS sample sizes across ancestries.**

PIPs for 100 simulated causal genes (A), the distribution of 90% credible gene set sizes for 100 simulated gene regions (B), and the sensitivity (C) from the single-ancestry FOCUS for EUR and AFR, MA-FOCUS, and baseline approach, varying complex trait variation explained by causal gene expression ($h_{GE}^{2}$) for each ancestry. The GWAS sample size was fixed at 503,717 and 13,313 for EUR and AFR, respectively. The eQTL panel sample size was fixed at 373 and 441 for EUR and AFR, respectively. See Methods section for default parameters. The black dashed lines indicate 90%. Error bars are constructed using a 95% confidence interval.

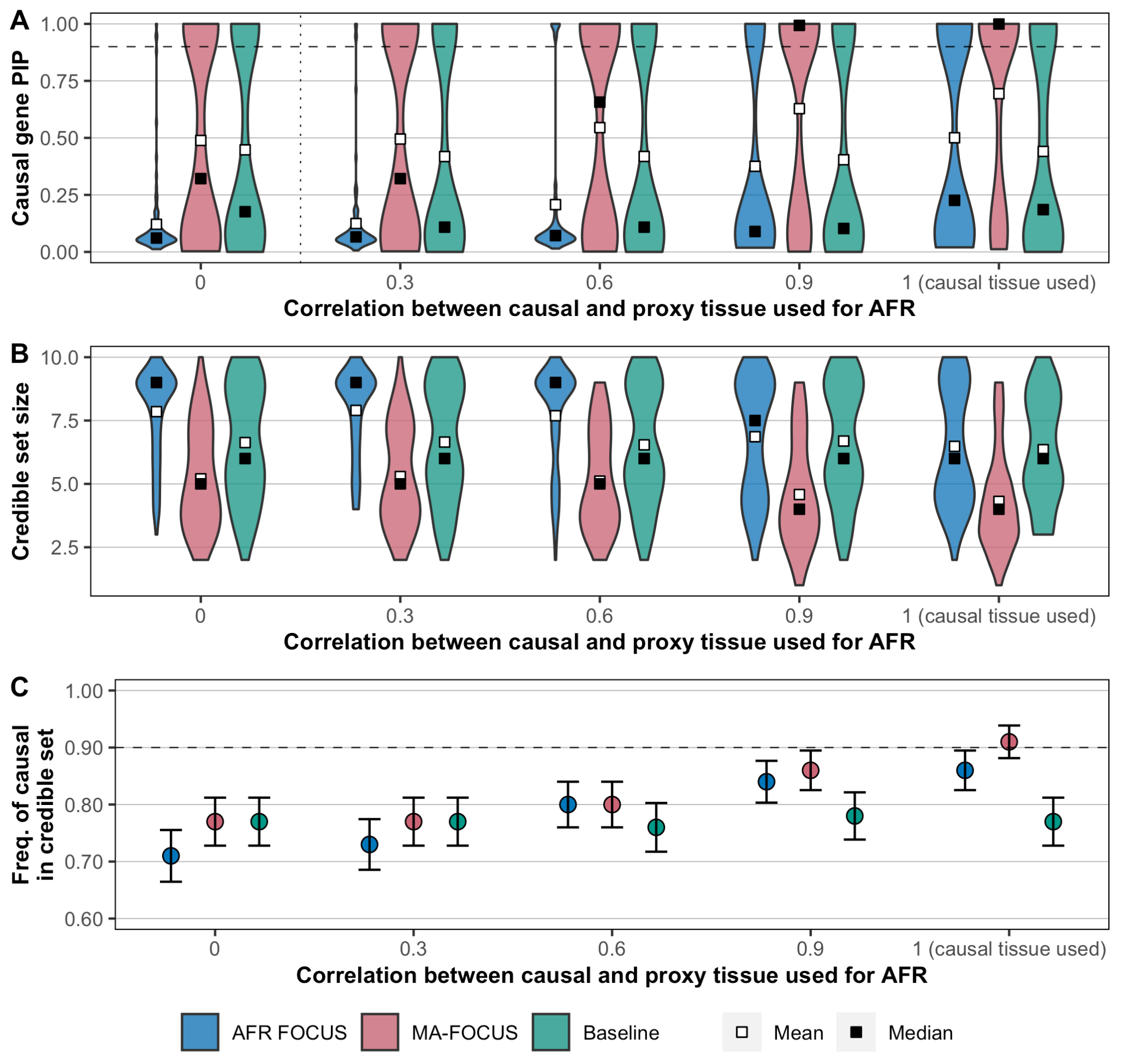

##### **Figure S10. MA-FOCUS outperforms the baseline method when proxy tissue is used.**

PIPs for 100 simulated causal genes (A), the distribution of 90% credible gene set sizes for 100 simulated gene regions (B), and the sensitivity (C) from the single-ancestry FOCUS for AFR, MA-FOCUS, and baseline approach, varying gene expression correlation between causal and proxy tissues. See Methods section for default parameters. The black dashed lines indicate 90%. Error bars are constructed using a 95% confidence interval.

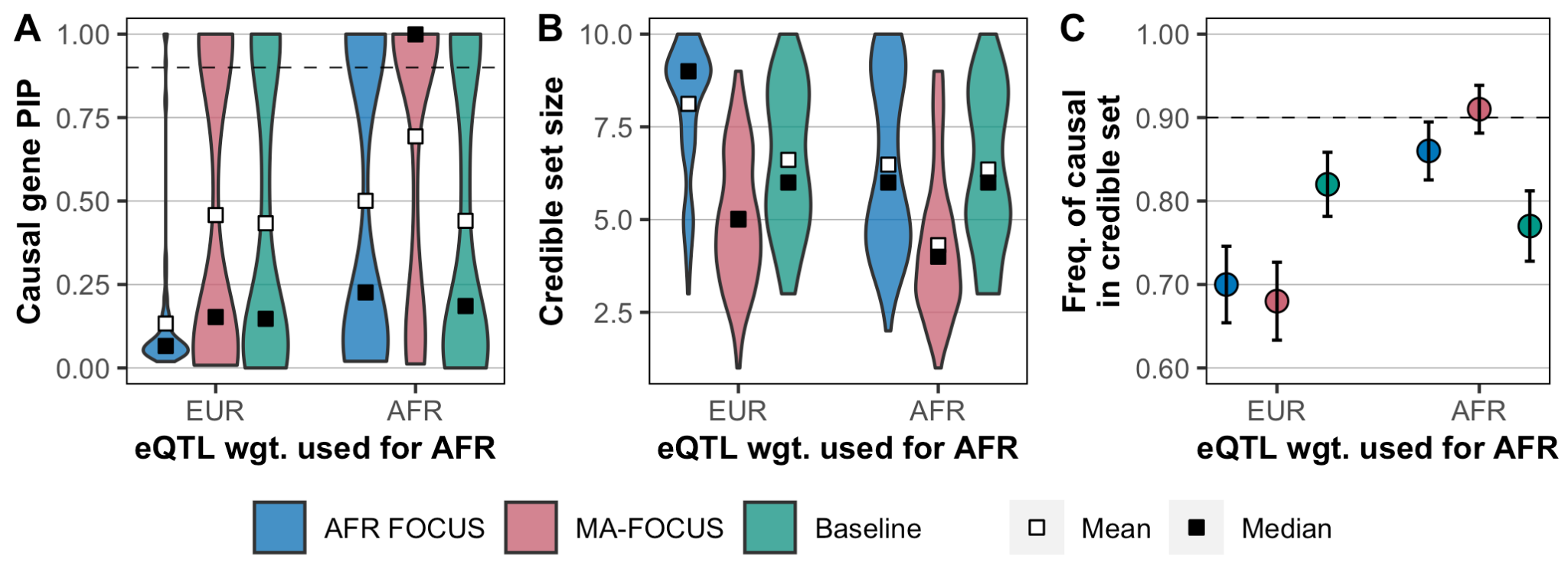

##### **Figure S11. MA-FOCUS remains robust when the EUR eQTL panel is substituted for AFR.**

PIPs for 100 simulated causal genes (A), the distribution of 90% credible gene set sizes for 100 simulated gene regions (B), and the sensitivity (C) from the single-ancestry FOCUS for AFR, MA-FOCUS, and baseline approach when EUR eQTL weights are substituted for AFR eQTL weights or still used original one. See Methods section for default parameters. The black dashed lines indicate 90%. Error bars are constructed using a 95% confidence interval.

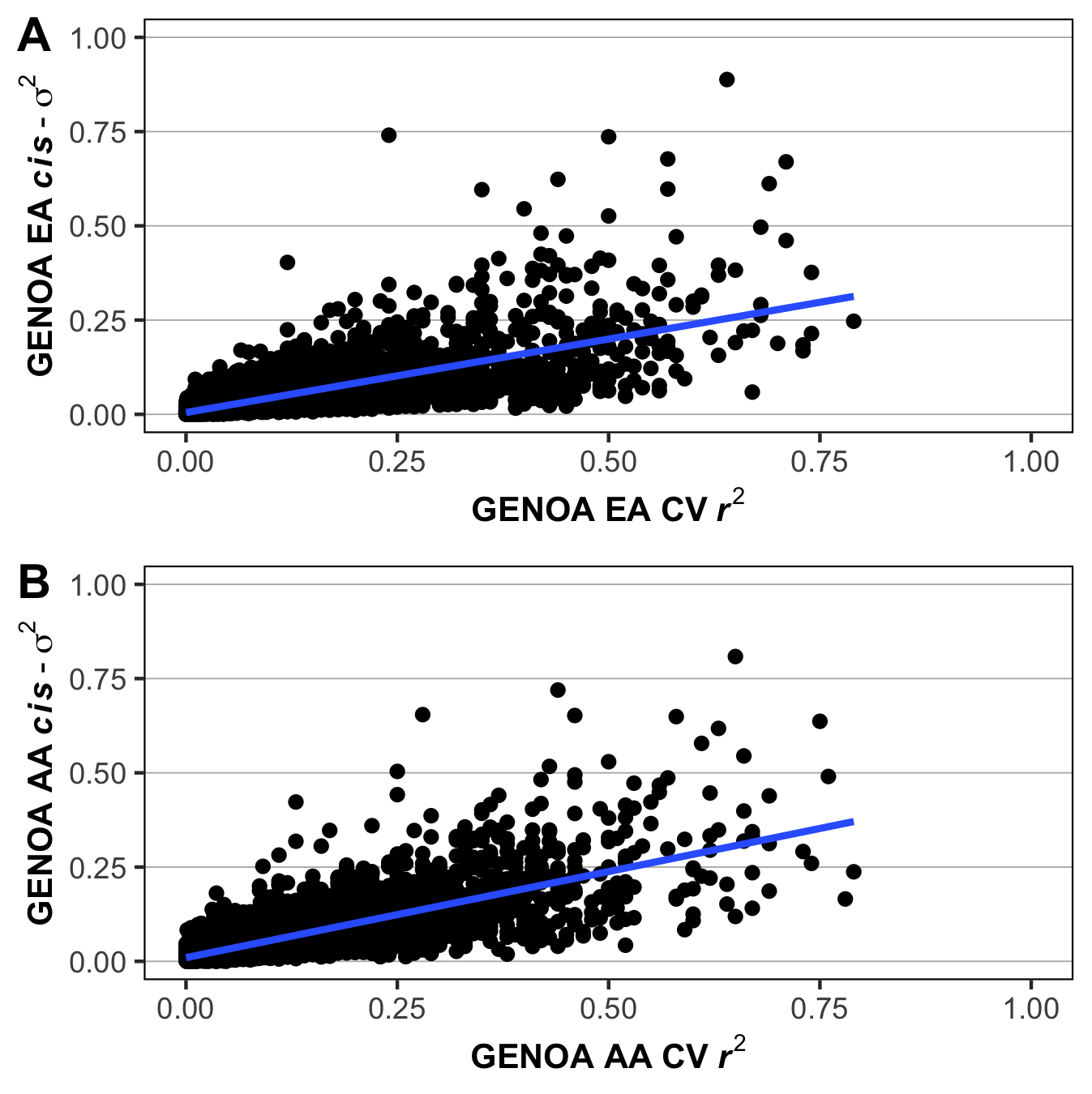

##### **Figure S12. Significant correlation between Cross-validation** $\boldsymbol{r}^{\boldsymbol{2}}$ **and genetic variance *cis-***$\boldsymbol{\sigma}_{\boldsymbol{g}}^{\boldsymbol{2}}$**indicates the better performance of the predictive models in more heritable genes.**

(A) (B). Each point represents a gene. The y-axis is *cis*-genetic variance *cis*-$\sigma_{g}^{2}$ estimated using GREML. The x-axis is the average cross-validation (CV) $r^{2}$ of the corresponding fitting model. The blue line is estimated using ordinary linear regression.

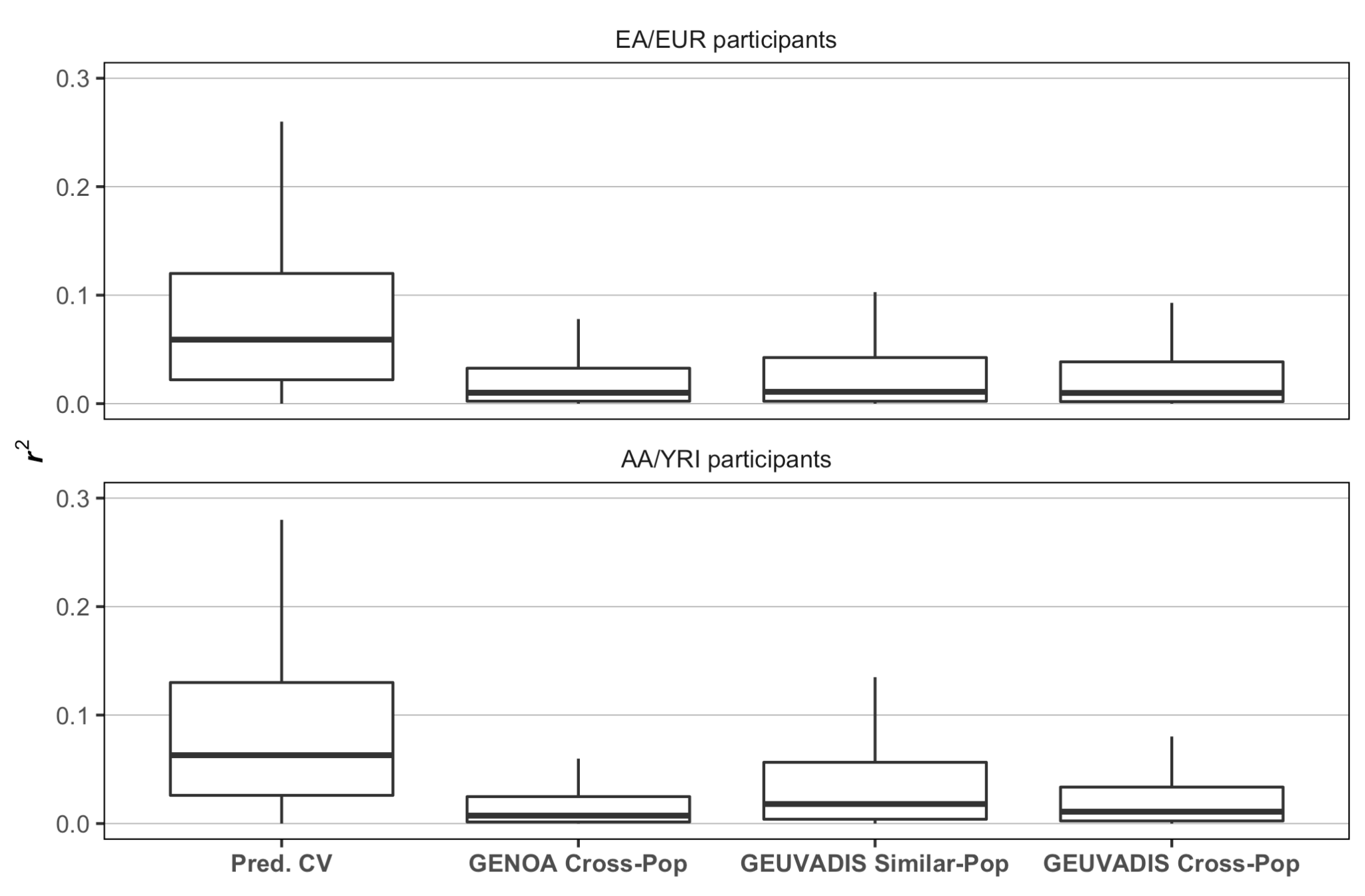

##### **Figure S13. GENOA-based prediction models replicate well in GEUVADIS individuals.**

We compared transportability of genetically predicted gene expression across various settings. The left-most boxplot (Pred. CV) is the in-sample cross-validation $r^{2}$ when fitting prediction models in GENOA. The next three reflect the distribution of $r^{2}$ computed using measured and predicted gene expression levels, when the source of the prediction model changes. The second reflects GENOA cross-ancestry weights (i.e. within GENOA EA->AA, or AA->EA). The third reflects GEUVADIS similar-ancestry weights (i.e. GENOA EA -> GEUVADIS EUR and GENOA AA -> GEUVADIS YRI). The final and fourth reflects GEUVADIS cross-ancestry weights (i.e. GENOA EA -> GEUVADIS YRI or GENOA AA -> GEUVADIS EUR).

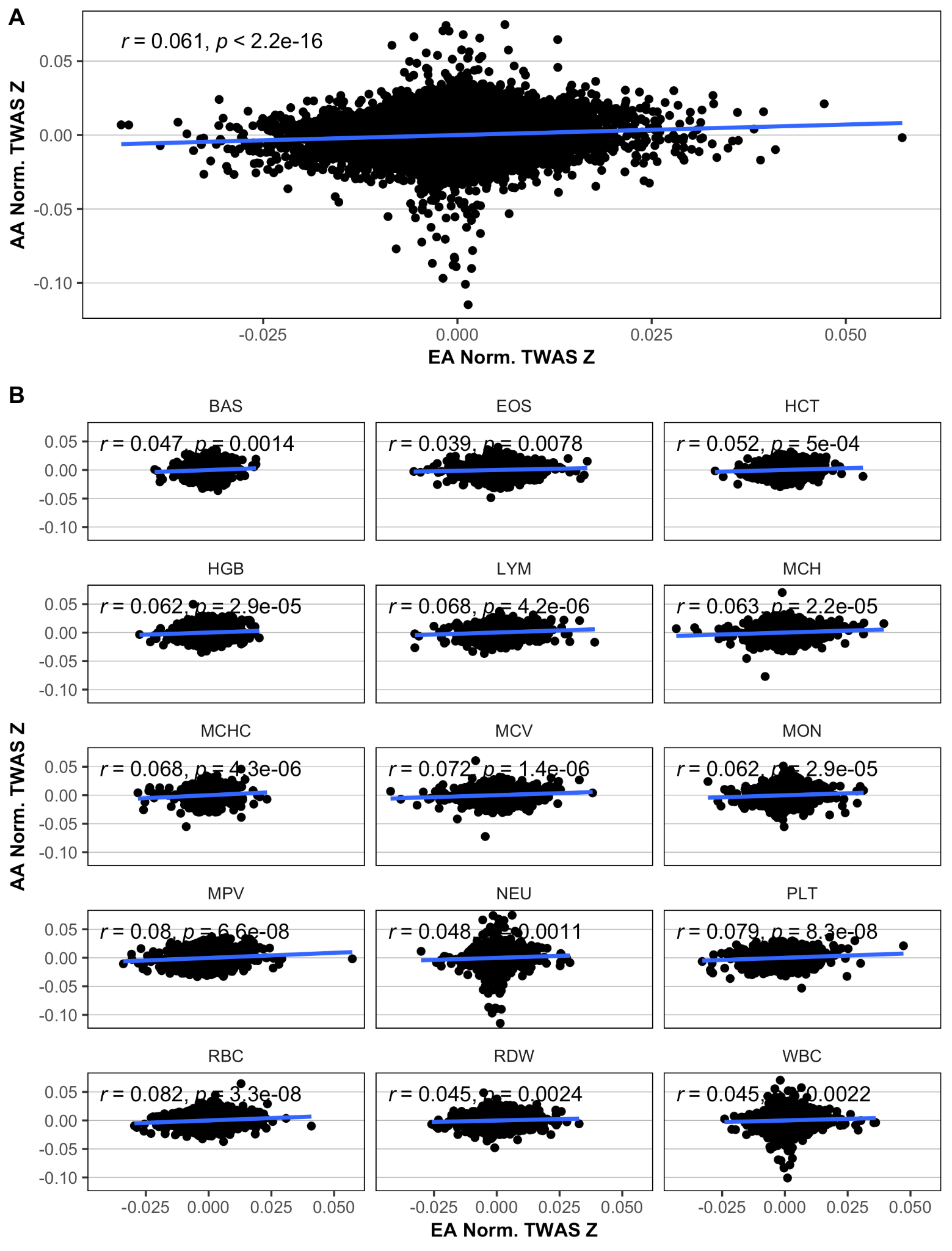

##### **Figure S14. Scatterplot of sample-size normalized TWAS z scores between EA and AA.**

(A) (B) Each point represents a gene. The blue line is a regression slope estimated using ordinary linear regression. Pearson’s method is used for the correlation. (A) is the integration across all blood traits and (B) is trait-specific break-down.

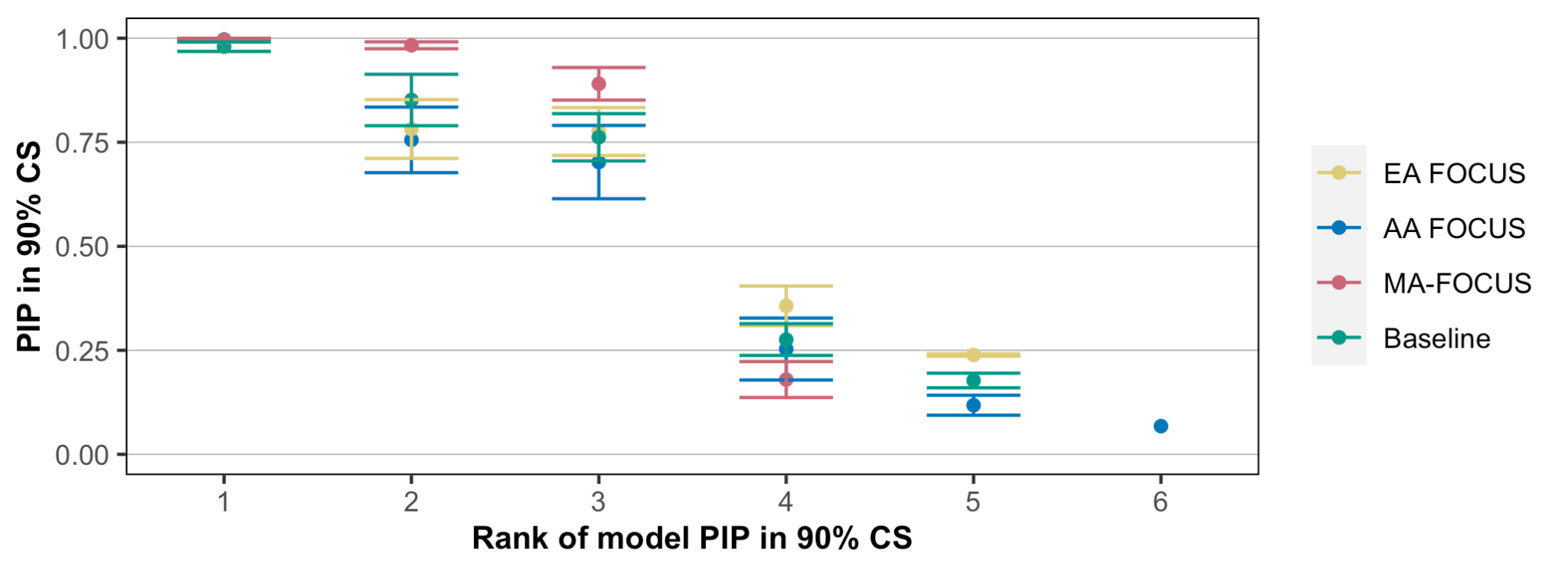

**Figure S15. Mean PIPs within ranked genes in credible sets by each method across all traits.**

The y-axis is the mean PIP of all models (including genes and null model) with corresponding rank in their respective credible sets by each method. The x-axis is the rank of the model. Error bars are constructed using a 95% confidence interval.

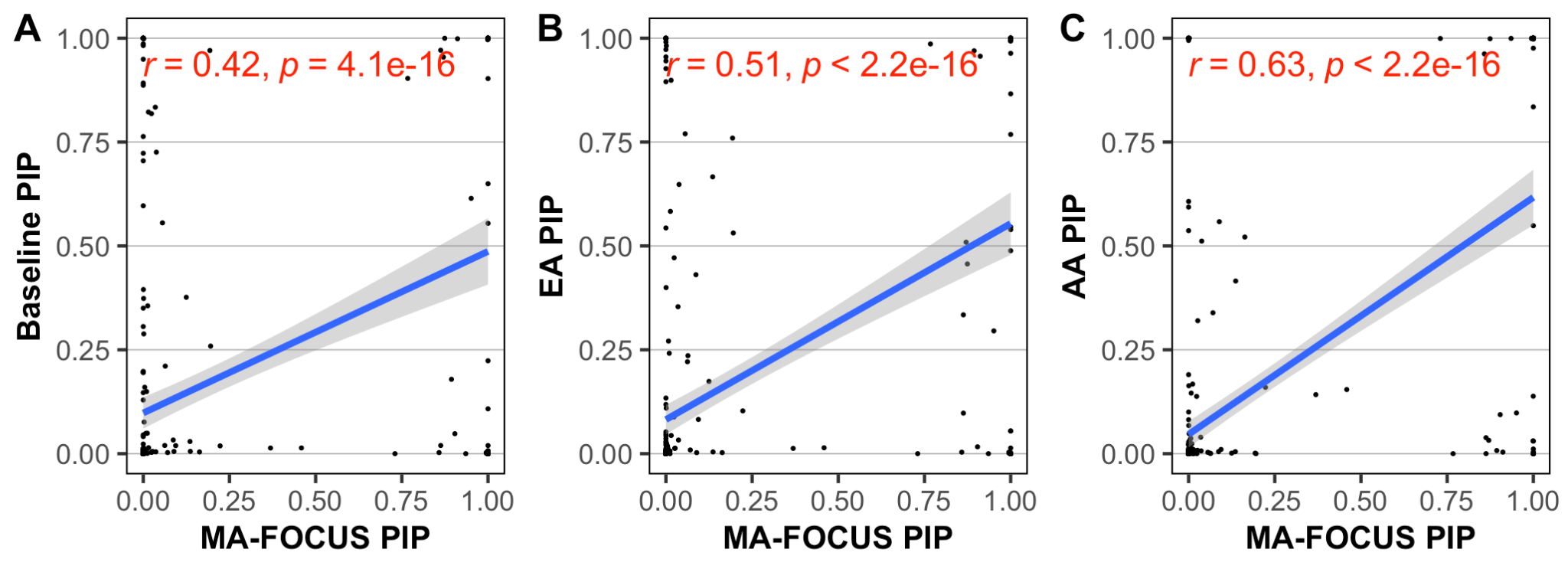

##### **Figure S16. PIPs are highly correlated between MA-FOCUS and other approaches.**

Each point represents a gene fine-mapped by each method. The blue line is estimated using ordinary linear regression. Pearson’s method is used for the correlation.

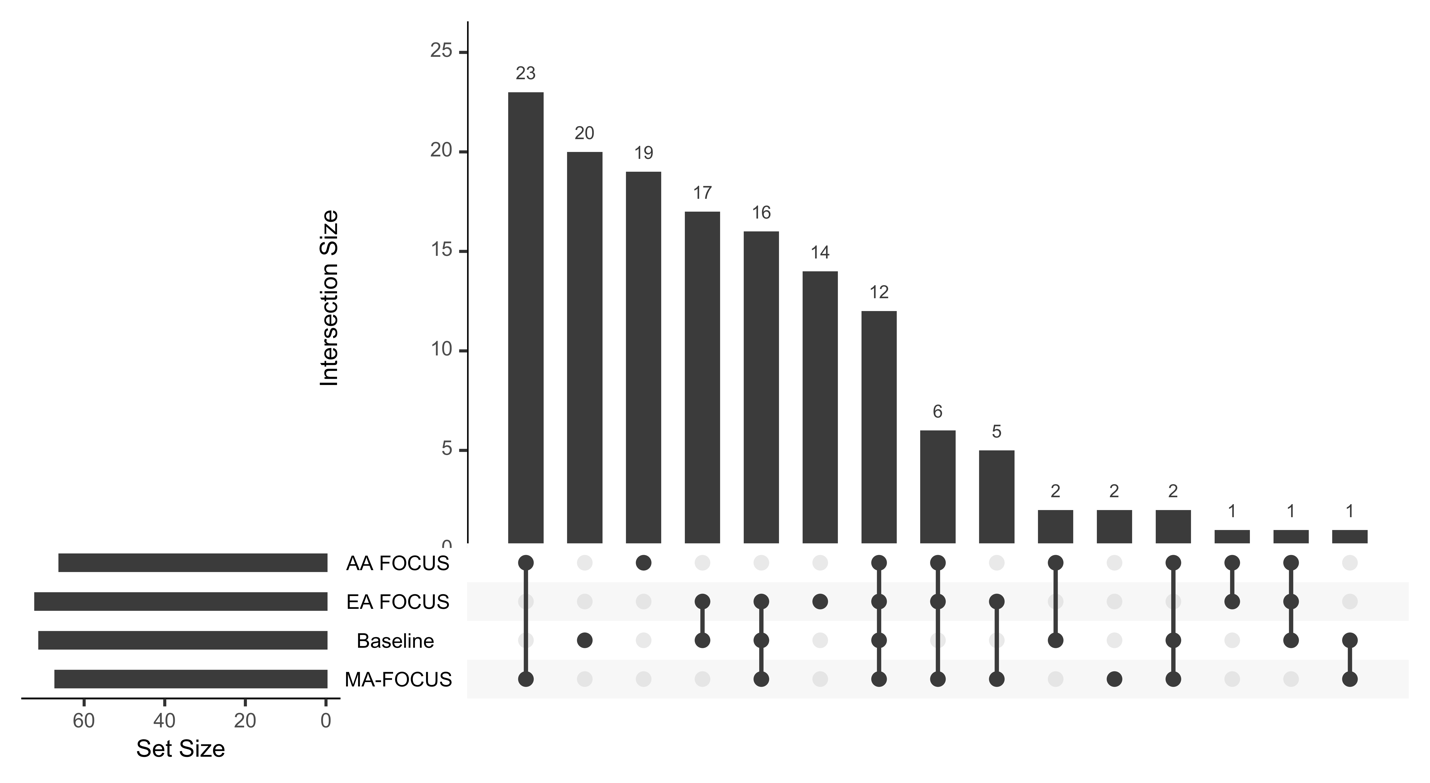

##### **Figure S17. Intersections of credible sets computed by MA-FOCUS, Baseline, and EA FOCUS.**

Results are calculated based on 22 regions that contain both EA and AA TWAS signals across all 10 traits. The UpsetR package is used to generate this plot.

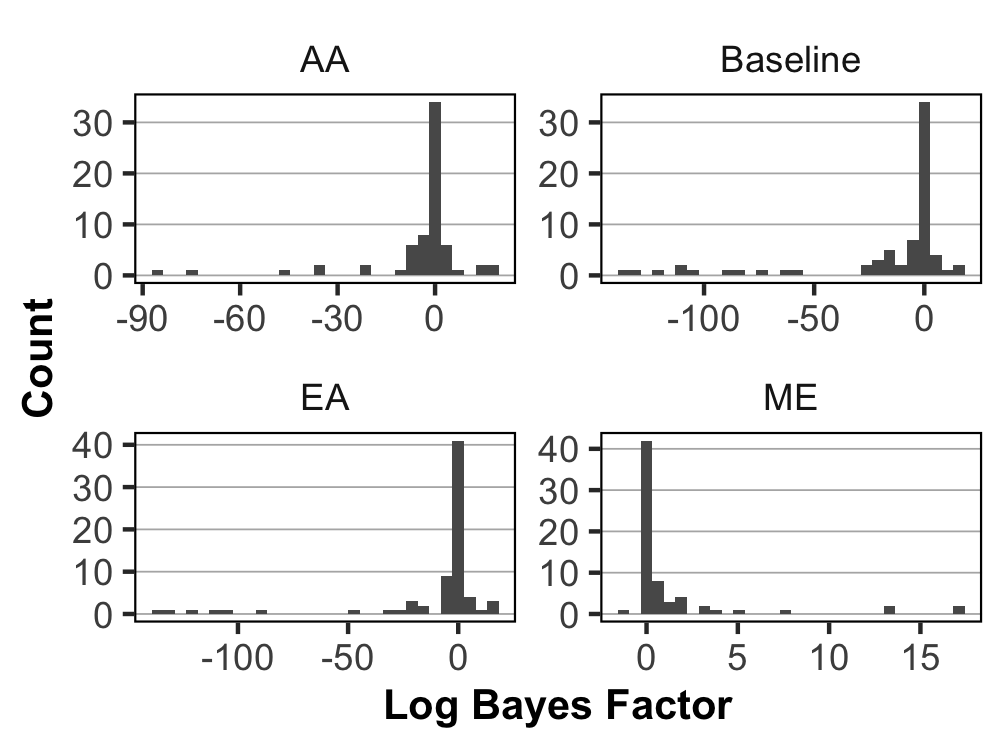

##### **Figure S18. Positive log-scale Bayes factors by MA-FOCUS indicate evidence for shared causal genes across ancestries.**

Bar plots for log Bayes factor where filtering on genes that are included in AA, Baseline, EA, and ME credible sets. Results are calculated based on 22 regions that contain both EA and AA TWAS signals across all 10 traits. See Methods section for how to compute the log-scale Bayes factors.

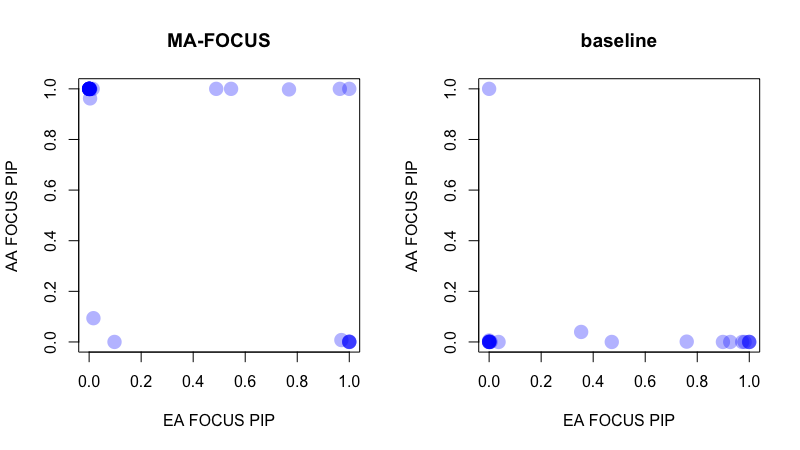

##### **Figure S19. Ancestry-specific fine-mapping PIPs for MA-FOCUS-specific and baseline-specific genes.**

MA-FOCUS-specific genes (i.e. genes that MA-FOCUS identified but the baseline approach did not) tend to have moderate high PIPs in either or both AA- or EA-specific FOCUS fine-mapping compared to baseline-specific genes (i.e. genes that the baseline approach identified but that baseline approach did not), which tend to show low PIPs in ancestry-specific fine-mapping. This suggests that, broadly, MA-FOCUS is better able to identify genes that have evidence of causality in at least one ancestry, while the baseline approach identifies genes that have weak or no evidence of causality in either ancestry.

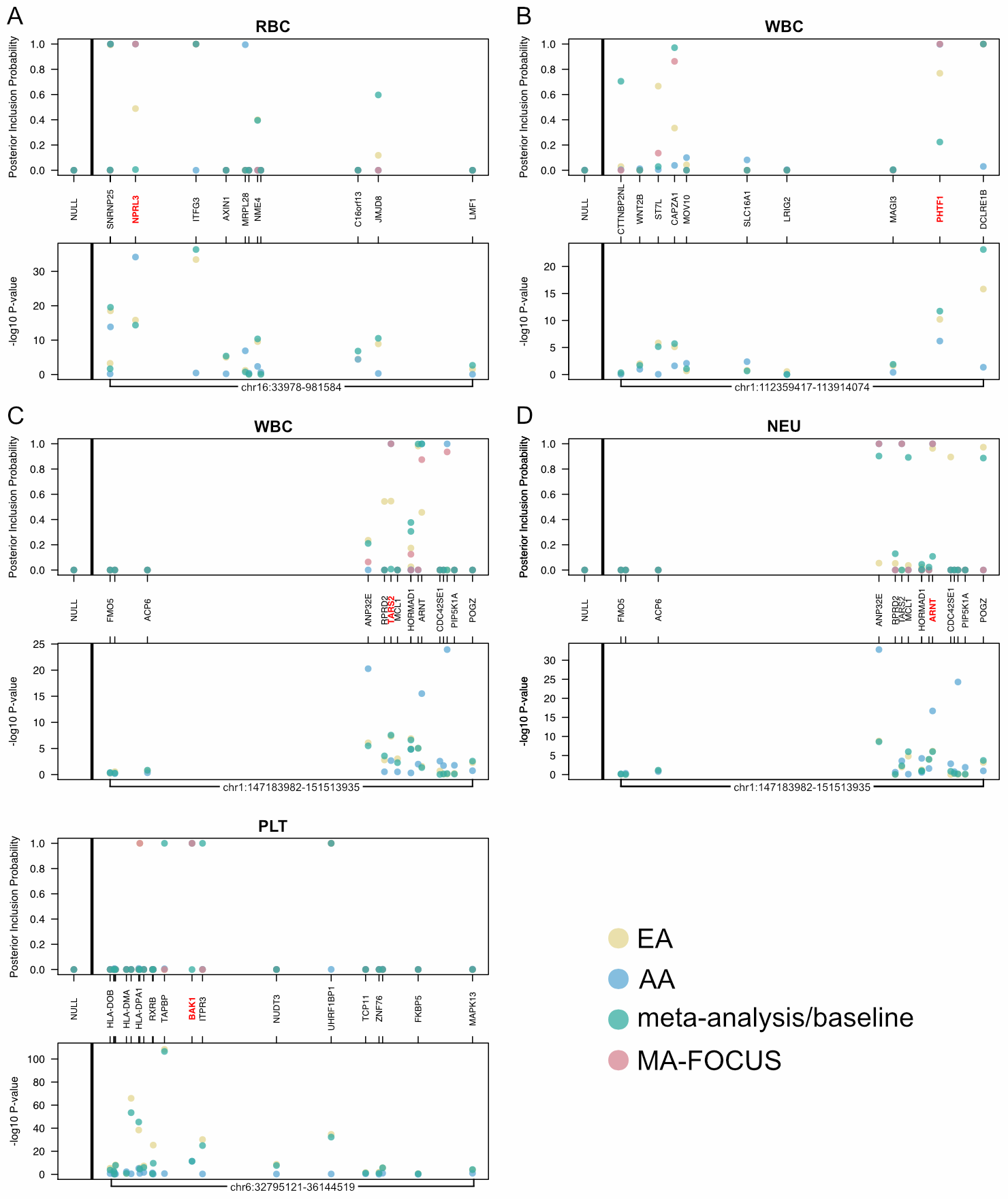

##### **Figure S20. PIPs and TWAS P-values at independent genomic regions with MA-FOCUS-specific genes.**

Each panel highlights a particular gene (red) in its genomic context; the top panels plot fine-mapping PIPs while the bottom panels plot TWAS P-values along the genome. The leftmost section of each top panel shows the PIP for the null model under each method.
